# The TRAPP complex mediates secretion arrest induced by stress granule assembly

**DOI:** 10.1101/528380

**Authors:** Francesca Zappa, Cathal Wilson, Giuseppe Di Tullio, Michele Santoro, Piero Pucci, Maria Monti, Davide D’Amico, Sandra Pisonero Vaquero, Rossella De Cegli, Alessia Romano, Moin A. Saleem, Elena Polishchuk, Mario Failli, Laura Giaquinto, Maria Antonietta De Matteis

## Abstract

The TRAnsport-Protein-Particle (TRAPP) complex controls multiple membrane trafficking steps and is thus strategically positioned to mediate cell adaptation to diverse environmental conditions, including acute stress. We have identified TRAPP as a key component of a branch of the integrated stress response that impinges on the early secretory pathway. TRAPP associates with and drives the recruitment of the COPII coat to stress granules (SGs) leading to vesiculation of the Golgi complex and an arrest of ER export. Interestingly, the relocation of TRAPP and COPII to SGs only occurs in actively proliferating cells and is CDK1/2-dependent. We show that CDK1/2 activity controls the COPII cycle at ER exit sites (ERES) and that its inhibition prevents TRAPP/COPII relocation to SGs by stabilizing them at the ERES. Importantly, TRAPP is not just a passive constituent of SGs but controls their maturation since SGs that assemble in TRAPP-depleted cells are smaller and are no longer able to recruit RACK1 and Raptor, rendering the cells more prone to undergo apoptosis upon stress exposure.

## Introduction

The TRAPP (TRAnsport Protein Particle) complex is a conserved multimolecular complex intervening in multiple segments of membrane trafficking along the secretory, the endocytic and the autophagy pathways (Kim et al., 2016).

Originally identified in yeast as a tethering factor acting in ER-to-Golgi trafficking, it was subsequently discovered to act as a GEF for Ypt1 and Ypt31/32 in yeast and for Rab1 and possibly Rab11 in mammals (Westlake et al., 2011; Yamasaki et al., 2009; Zou et al., 2012). The TRAPP complex has a modular composition and is present as two forms in mammals: TRAPPII and TRAPPIII which share a common heptameric core (TRAPPC1, TRAPPC2, TRAPPC2L, TRAPPC3, TRAPPC4, TRAPPC5, TRAPPC6) and additional subunits specific for TRAPPII (TRAPPC9/ TRAPPC10) or for TRAPPIII (TRAPPC8, TRAPPC11, TRAPPC12, TRAPPC13) (Sacher et al., 2018). TRAPPII has been implicated in late Golgi trafficking while TRAPPIII has a conserved role in the early secretory pathway (ER-to-Golgi) and in autophagy (Scrivens et al., 2011; Yamasaki et al., 2009). Recent lines of evidence have expanded the range of activities of the TRAPP complex by showing that it takes part in a cell survival response triggered by agents that disrupt the Golgi complex (Ramírez-Peinado et al., 2017) and can drive the assembly of lipid droplets in response to lipid load (Li et al., 2017).

The importance of the TRAPP complex in humans is testified by the deleterious consequences caused by mutations in genes encoding distinct TRAPP subunits. Mutations in TRAPPC2L, TRAPPC6A, TRAPPC6B, TRAPPC9 and TRAPPC12 cause neurodevelopmental disorders leading to intellectual disability and dysmorphic syndromes (Harripaul et al., 2018; Khattak and Mir, 2014; Milev et al., 2018, 2017, p. 12; Mohamoud et al., 2018), mutations in TRAPPC11 lead to ataxia (Koehler et al., 2017), and mutations in TRAPPC2 lead to the spondyloepiphyseal dysplasia tarda (SEDT) (Gedeon et al., 1999).

SEDT is characterized by short stature, platyspondyly, barrel chest and premature osteoarthritis that manifest in late childhood/prepubertal age. We have shown that pathogenic mutations or deletion of TRAPPC2 alter the ER export of procollagen (both type I and type II) and that TRAPPC2 interacts with the procollagen escort protein TANGO1 and regulates the cycle of the GTPase Sar1 at the ER exit sites (ERES) (Venditti et al., 2012)). The Sar1 cycle in turn drives the cycle of the COPII coat complex, which mediates the formation of carriers containing neo-synthesized cargo to be transported to the Golgi complex. While this role of TRAPPC2 in the ER export of PC might explain the altered ECM deposition observed in patient’s cartilage (Venditti et al., 2012; Tiller et al., 2001), it leaves the late onset of the disease signs unexplained. We hypothesized that the latter could be due to an inability to maintain long-term cartilage tissue homeostasis, possibly due to an impaired capacity of chondrocytes to face the physiological stresses that underlie and guide their development and growth. These include mechanical stress that can induce oxidative stress leading to apoptosis (Henrotin et al., 2003; Zuscik et al., 2008).

Here, by analyzing the cell response to different stresses, we show that the TRAPPC2 and the entire TRAPP complex, are components of stress granules (SGs), membrane-less organelles that assemble in response to stress (Protter and Parker, 2016).

The recruitment of TRAPP to SGs has multiple impacts on the stress response as it induces the sequestration of Sec23/Sec24 (the inner layer of the COPII complex) onto SGs thus inhibiting trafficking along the early secretory pathway, leads to the inactivation of the small GTPase Rab1 with a consequent disorganization of the Golgi complex, and is required for the recruitment of signalling proteins, such as Raptor and RACK1, to SGs thus contributing to the anti-apoptotic role of SGs. Interestingly, TRAPP and COPII recruitment to SGs only occurs in actively proliferating cells and is under control of cyclin-dependent-kinases (CDK 1 and 2). The TRAPP complex thus emerges as a key element in conferring an unanticipated plasticity to SGs, which adapt their composition and function to cell growth activity, reflecting the ability of cells to adjust their stress response to their proliferation state and energy demands.

## Results

### TRAPP redistributes to SGs in response to different stress stimuli

To investigate the involvement of TRAPPC2 in the stress response, we exposed chondrocytes (Figure 1A) or HeLa cells (Figure 1B) to sodium arsenite (SA) treatment that induces oxidative stress. We have shown previously that TRAPPC2 associate with ERES under steady state conditions (Venditti et al., 2012). SA treatment led to dissociation of TRAPPC2 from ERES and a complete relocation to roundish structures. Oxidative stress is known to lead to the formation of stress granules (SGs; Anderson and Kedersha, 2002), therefore, we considered the possibility that these TRAPPC2-positive structures might be SGs. Indeed, colabeling with an anti-eIF3 antibody, a canonical SG marker (Aulas et al., 2017), showed the co-localization of TRAPPC2 with eIF3 after SA treatment (Figure 1A,B). Other TRAPP components, such as TRAPPC1 and TRAPPC3, exhibited the same response to SA treatment (Figure 1C). TRAPP recruitment to SGs also occurs in response to heat shock (Figure 1—figure supplement 1A), and in different cell lines (Figure 1—figure supplement 1B,C). In addition, TRAPP elements associate to SGs in yeast cells exposed to heat stress (Figure 1—figure supplement 2, Figure 1—supplement video 1 and 2) thus suggesting that this is a conserved process in evolution. Of note, the association of TRAPP with SGs is fully reversible after stress removal (Figure 1—figure supplement 3).

**Figure 1.**
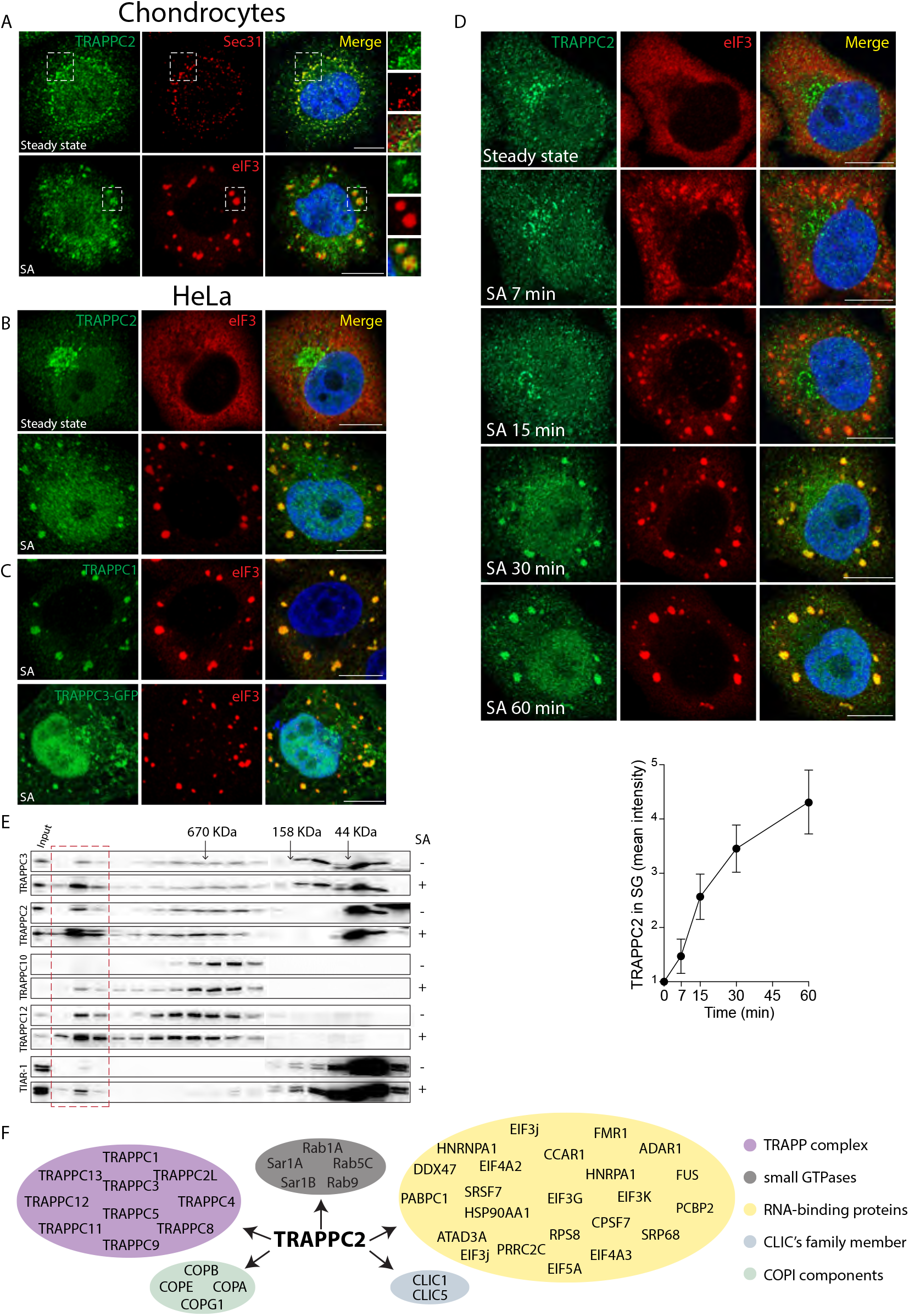
The TRAPP complex is recruited to stress granules. **(A,B)** Localization of endogenous TRAPPC2 under steady state conditions or after sodium arsenite (SA) treatment in (A) a rat chondrosarcoma cell line (300 μM, 1 hr) and (B) in HeLa cells (300 μM, 30 min). Fluorescence microscopy of fixed cells using antibodies against TRAPPC2, eIF3 (to label SGs), Sec31 (to label ERES). DAPI (blue). Small panels in (A) show details and merge of boxed areas. **(C)** Localization of endogenous TRAPPC1 and of GFP-TRAPPC3 to eIF3-labeled SGs in HeLa cells after SA treatment. **(D)** Representative images of a time course analysis of TRAPPC2 redistribution to SGs. Cells were treated as in (B). Graph, quantification of TRAPPC2 localization at SGs over time (the ratio between TRAPPC2 (mean fluorescence intensity) in SG puncta and cytosolic TRAPPC2). Mean ± s.e.m. n = 50 cells per experiment, N = 3. Scale bar, 10 μm in (A-D) **(E)** Extracts from untreated and SA-treated HeLa cells were fractionated by size exclusion chromatography on a Superose6 column. Fractions were analyzed by Western blot using antibodies against TRAPPC2 and TRAPPC3 (components of both TRAPPII and TRAPPIII complexes), TRAPPC10 (a component of the TRAPPII complex), TRAPPC12 (a component of the TRAPPIII complex), and against the SG marker TIAR-1. TRAPP components co-fractionated with a high molecular weight complex (red dashed box) containing TIAR-1 in SA-treated cells. Arrows indicate protein standards: thyroglobulin (670 kDa), *g*-globulin (158 kDa), ovalbumin(44 kDa). **(F)** Schematic representation of MS/MS analysis of TRAPPC2 interactors. 251 out of 512 (46%) proteins are RNA-binding proteins, including several SG proteins (yellow ellipse). **Figure 1—figure supplement 1**. Re-localization of TRAPP to SGs is independent of stress stimulus and cell type. **Figure 1—figure supplement 2**. Stress-induced localization of Bet3 (TRAPPC3) to SGs in yeast cells. **Figure 1—supplement video 1**. Time lapse microscopy of Bet3 (TRAPPC3) and Pab1 at 30°C in yeast cells. **Figure 1—supplement video 2**. Time lapse microscopy of Bet3 (TRAPPC3) and Pab1 at 46°C in yeast cells. **Figure 1—figure supplement 3**. Association of TRAPP with SGs is reversible after SG disassembly.

We studied the kinetics of TRAPPC2 redistribution to SGs. The appearance of eIF3-positive SGs occurred 7 min after SA treatment (Figure 1D), while TRAPPC2 relocalization to SGs began 15 min after exposure to stress and gradually increased over time, with massive recruitment occurring 60 min after treatment, thus lagging behind the initial assembly of SGs (Figure 1D).

We observed that oxidative stress did not affect the overall integrity of the TRAPP complexes (both TRAPPII and TRAPPIII), as analyzed by gel filtration. However, in stressed cells TRAPPII and TRAPPIII components were also found in higher molecular weight fractions (Figure 1E) containing the SG component TIAR-1 (Figure 1E), possibly reflecting the SG-associated TRAPP.

To investigate the mechanisms that could mediate the recruitment of TRAPP to SGs, we performed a proteomics analysis of the TRAPPC2 interactors (see Materials and methods). This analysis (Figure 1F, Table S1), confirmed known TRAPPC2 interactors (e.g. TRAPP complex components, Rab1, COPII and COPI) and revealed the presence of many RNA binding proteins (RBPs), including those with a central role in the assembly of SGs (Jain et al., 2016). This suggested that, as described for other components of SGs, TRAPPC2 may be recruited to growing SGs by a piggyback mechanism (Anderson and Kedersha, 2008), i.e. via the interaction with RBPs that are components of SGs.

Altogether, these results led us to conclude that following stress the TRAPP complex, through TRAPPC2 interaction with RBPs, becomes a SG component.

### TRAPPC2 is required for recruiting COPII to SGs

The translocation of TRAPP to SGs prompted us to investigate whether other components of membrane trafficking machineries behaved similarly. We screened different coat complex components (COPI, COPII, clathrin adaptors, clathrin), and other cytosolic proteins associated with the exocytic and endocytic pathways. Out of the 15 proteins tested, only components of the inner layer of the COPII coat, Sec24 (Figure 2A, B) and Sec23 (Figure 2—figure supplement 1A), but none of the others (Figure 2C, and Figure 2—figure supplement 1B-N, Figure 2— figure supplement 2) associated with SGs. Notably, Sec24 was recruited to SGs with kinetics similar to those of the TRAPP complex (Figure 2B).

**Figure 2.**
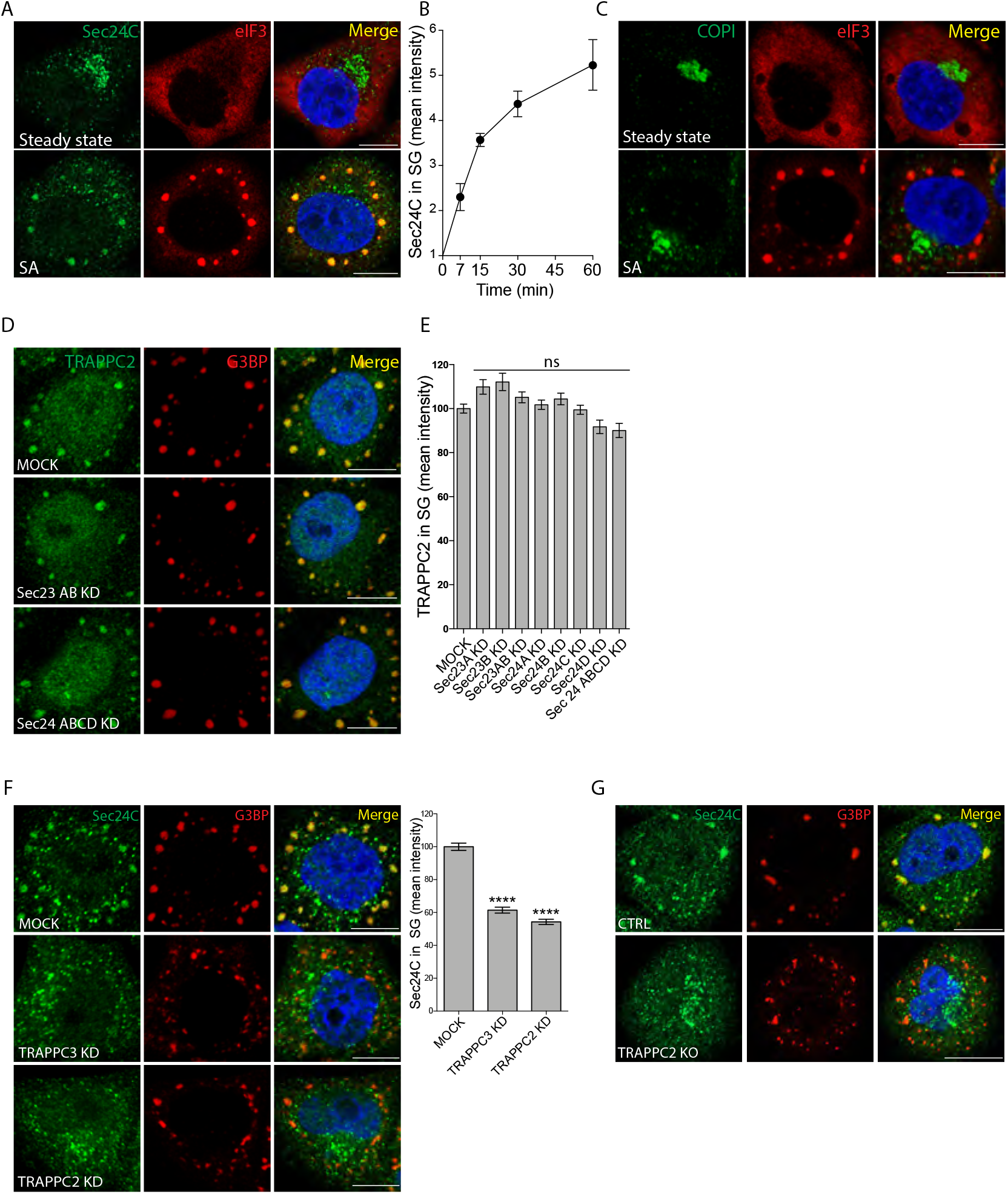
TRAPPC2 is required for Sec23/24 relocalization to SGs. **(A)** HeLa cells, untreated or treated with SA (300 μM, 30 min), were fixed and visualized by fluorescence microscopy using anti-Sec24C Ab, anti-eIF3 Ab, and DAPI (blue). **(B)** Quantification of Sec24C redistribution to SGs over time after SA treatment (the ratio between Sec24C (mean fluorescence intensity) in SG puncta and cytosolic Sec24C). Mean ± s.e.m. n = 50 cells per experiment, N = 3. **(C)** Cells treated as in (A) stained with an antibody recognizing the coatomer I (COPI), which does not relocalize to SGs (see also Figure 2—figure supplement 1A-N). **(D)** Representative images of TRAPPC2 localization in Sec23AB-KD and Sec24ABCD-KD cells treated with SA. Cells were fixed and visualized by fluorescence microscopy using anti-TRAPPC2 Ab, anti-G3BP Ab (to label SGs) and DAPI (blue). **(E)** Quantification of TRAPPC2 redistribution to SGs after KD of the indicated Sec23 and Sec24 combinations, calculated as the ratio between TRAPPC2 (mean intensity) in SG puncta and cytosolic TRAPPC2. Mean ± s.e.m., n = 40-60 cells per experiment, N = 3. ns: not significant. **(F)** Representative images of Sec24C localization at SGs (stained for G3BP) in TRAPPC3-KD and TRAPPC2-KD cells treated with SA. Depletion of the entire TRAPP complex (via TRAPPC3 depletion) or of only TRAPPC2 reduces Sec24C recruitment. Graph, quantification of Sec24C at SGs, calculated as in (A). Mean ± s.e.m., n = 40-60 cells per experiment, N = 3. **** P < 0.0001. **(G)** Representative images of Sec24C localization at SGs (stained for G3BP) in TRAPPC2-KO cells treated with SA. Scale bar 10 μm. **Figure 2—figure supplement 1.** Sec24C and Sec23, but not components of other coat complexes or other cytosolic proteins associated with the exocytic and endocytic pathways, associate with SGs. **Figure 2—figure supplement 2.** Sec16 does not significantly associate with SGs nor is it required for the recruitment of COPII to SGs. **Figure 2—figure supplement 3.** Evaluation of Sec23, Sec24, TRAPPC2 and TRAPPC3 knock down efficiency.

Since TRAPP and COPII are known to interact and TRAPP is recruited to ERES in a Sar1- and COPII-dependent manner (Lord et al., 2011; Venditti et al., 2012), we asked whether COPII and TRAPP recruitment to SGs is interdependent. The depletion of the COPII inner layer proteins Sec23 and Sec24 had no impact on the recruitment of the TRAPP complex to SGs (Figure 2D,E, Figure 2—figure supplement 3A,B) while the depletion of the entire TRAPP complex (by KD of TRAPPC3, which destabilizes the entire complex) or of the TRAPPC2 subunit (by KD of TRAPPC2 that leaves unaltered the rest of the TRAPP complex; Venditti et al., 2012 or by TRAPPC2 KO via CRISPR-CAS9) abrogated the recruitment of COPII components to SGs (Figure 2F, G, Figure 2—figure supplement 3C). These results indicated that the TRAPP complex, through its component TRAPPC2, drives the recruitment of COPII to SGs formed in response to acute oxidative stress.

It is worth mentioning that COPII components have been reported to partition in membrane-less bodies called “Sec bodies” in Drosophila S2 cells (Zacharogianni et al., 2014). Sec bodies, which are distinct from SGs as they are devoid of ribonucleoproteins, form exclusively in response to prolonged aa starvation and not in response to acute oxidative stress (Zacharogianni et al., 2014). The formation of Sec bodies in mammalian cells has remained unclear (Zacharogianni et al., 2014). In fact, we found that COPII components (Sec24C and Sec31A) do not form Sec bodies as they remain associated with ERES in aa starved mammalian cells (Figure 2—figure supplement 2A, B). We also found that Sec16, which is required for the formation of Sec bodies and SGs in response to aa starvation in Drosophila S2 cells (Zacharogianni et al., 2014; Aguilera-Gomez et al., 2017), neither significantly associates with SGs formed in response to stress in mammalian cells (Figure 2—figure supplement 2C,D) nor is it required for the recruitment of COPII to these SGs (Figure 2—figure supplement 2E,F).

Altogether these results indicate that COPII components can undergo two distinct phase separation events: they can associate in a TRAPP-dependent fashion to SGs in response to acute stress in mammalian cells (this report) and they can assemble in a Sec16-dependent manner in “Sec bodies” in Drosphila S2 cells in response to aa starvation (Zacharogiani et al., 2014). Interestingly, these two distinct phase separation events lead to different functional consequences (see below).

We then moved to investigate the nature of the association of COPII to SGs since the COPII coat cycles in a very dynamic fashion at ERES (D’Arcangelo et al., 2013). To this end, we permeabilized living cells (Kapetanovich et al., 2005) at steady state and upon stress induction. Treatment with digitonin forms pores at the plasma membrane that allows the exit of the cytosolic content into the extracellular space. Under these conditions, we found that the COPII pool recruited to SGs is less dynamic than the pool cycling at the ERES, since COPII remained associated with SGs in permeabilized cells exposed to stress while the COPII pool cycling at the ERES was completely lost upon permeabilization of unstressed cells (Figure 3A). We then assessed the possible interplay between the COPII fraction at ERES and the pool that eventually relocates to SGs. We asked whether changes in the association of COPII at the ERES could have an impact on the recruitment of COPII to SGs. To this end, we stabilized COPII and TRAPP at ERES by expressing either a constitutively active GTP-bound Sar1 mutant (Sar1-H79G) or decreasing the rate of GTP hydrolysis on Sar1 by depleting Sec31, which is a co-GAP that potentiates by an order of magnitude the Sar1 GAP activity of Sec23-24 (Bi et al., 2007). Under both conditions, COPII and TRAPP were more tightly associated with ERES, while their translocation to SGs upon exposure to stress was significantly reduced (Figure 3B,C; Figure 3—figure supplement 1). Thus, COPII/TRAPP recruitment to SGs is commensurate with the rate of their association at the ERES indicating that the cytosolic pool of COPII generated through the fast cycling at ERES is the one available to be recruited to SGs upon stress.

**Figure 3.**
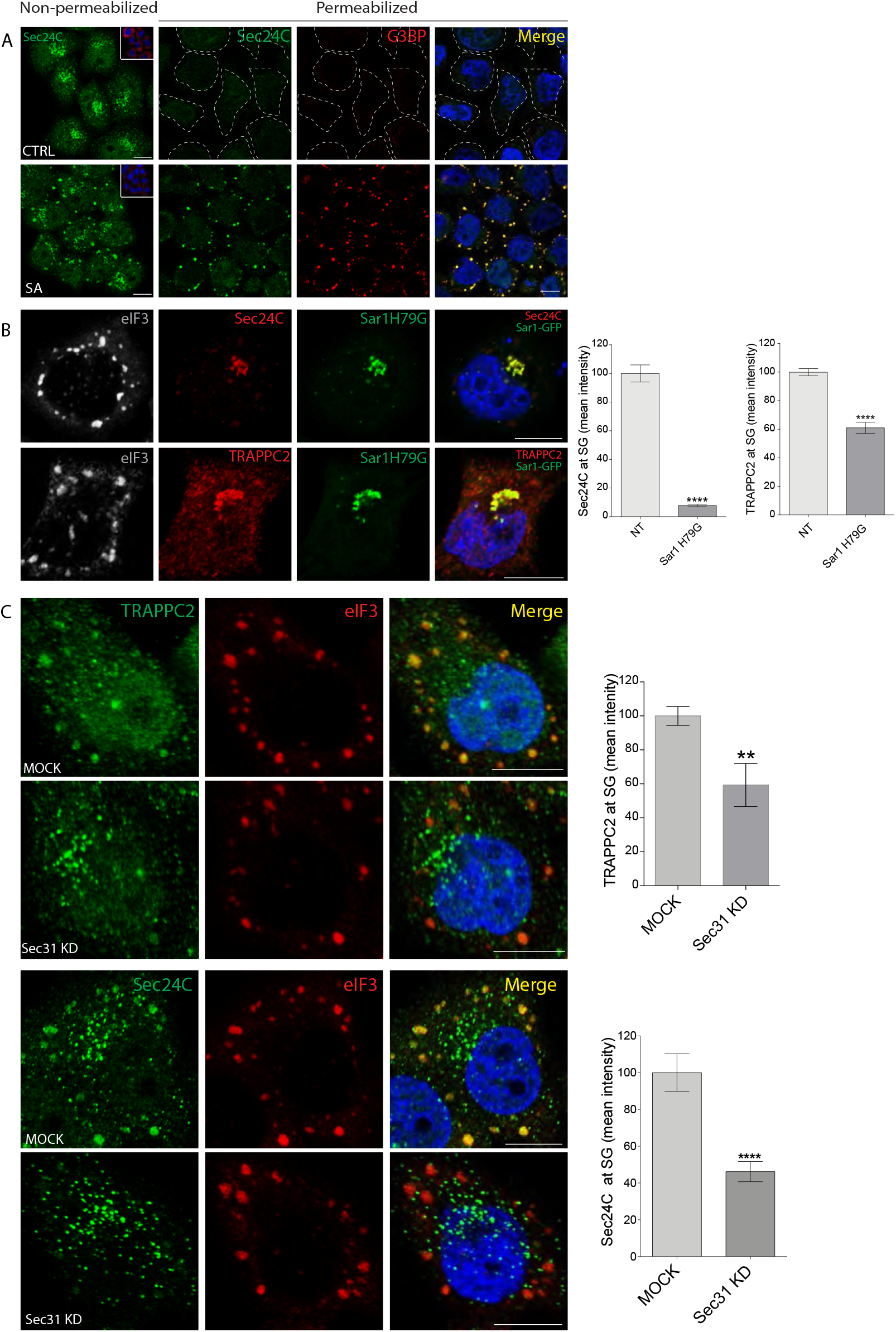
Comparison and relationships between the ERES-associated and the SG-associated pools of TRAPP and COPII. **(A)** TRAPP and COPII associate more stably with SGs than with ERES. The membrane association of Sec24C was evaluated in non-permeabilized or permeabilized cells with or without SA treatment, as indicated. G3BP was used as an SG marker. Left panel insets, G3BP staining in non-permeabilized cells. Dashed white lines show the outline of permeabilized SA-untreated cells. Blue, DAPI. **(B-C)** Stabilizing TRAPP and COPII at the ERES prevents their relocalization to SGs. **(B)** HeLa cells overexpressing GFP-Sar1H79G were treated with SA and immunostained for Sec24C, TRAPPC2, and eIF3, as indicated. Blue, DAPI. Graphs show quantification of Sec24C or TRAPPC2 at SGs in GFP-Sar1H79G-expressing cells relative to non-transfected cells. Data are the ratio between Sec24C or TRAPPC2 mean intensity in SG puncta and Sec24C or TRAPPC2 mean intensity at ERES, expressed as a percentage of the non-transfected cells. Mean ± s.e.m. n = 60-70, three independent experiments ****p<0.0001. **(C)** Effect of Sec31A depletion on TRAPPC2 and Sec24C recruitment to SGs. Cells were mock-treated or KD for Sec31, treated with SA (300 μM, 30 min), and then immunostained for TRAPPC2, Sec24C and eIF3 as indicated. Graph, quantification of Sec24C and TRAPPC2 at SGs, calculated as in (B). Mean ± s.e.m. of TRAPPC2 and Sec24C, one representative experiment; n=60-80. ** p<0.001, ****p<0.0001. Scale bar, 10 μm. **Figure 3—figure supplement 1.** Effect of Sec31 depletion on Sec24C localization at ERES.

### The association of TRAPP/COPII to SG is under the control of CDK1/2

To dissect the regulation of TRAPP/COPII recruitment to SGs, we screened a library of kinase inhibitors for their ability to affect the re-localization of COPII to SGs. HeLa cells were pre-treated with kinase inhibitors for 150 min and then exposed to SA for a further 30 min in the continuous presence of the inhibitors. Out of the 273 inhibitors tested, 194 had no effect on SG formation or on COPII recruitment, 13 were generally toxic (65% mortality) at the tested concentrations, one (AZD4547) affected the formation of SGs, while the remaining 32 specifically inhibited (by at least 35%) the association of Sec24C with SGs without affecting SG formation (Figure 4A). Enrichment analysis on this last group of inhibitors highlighted that CDK inhibitors were the most represented class (Figure 4B). Western blot analysis confirmed that CDK activity was impaired in cells treated with the CDK inhibitors (Figure 4C). In addition, we observed that there is an increase in CDK activity in SA-treated cells that was sensitive to CDK inhibitors (Figure 4C).

**Figure 4.**
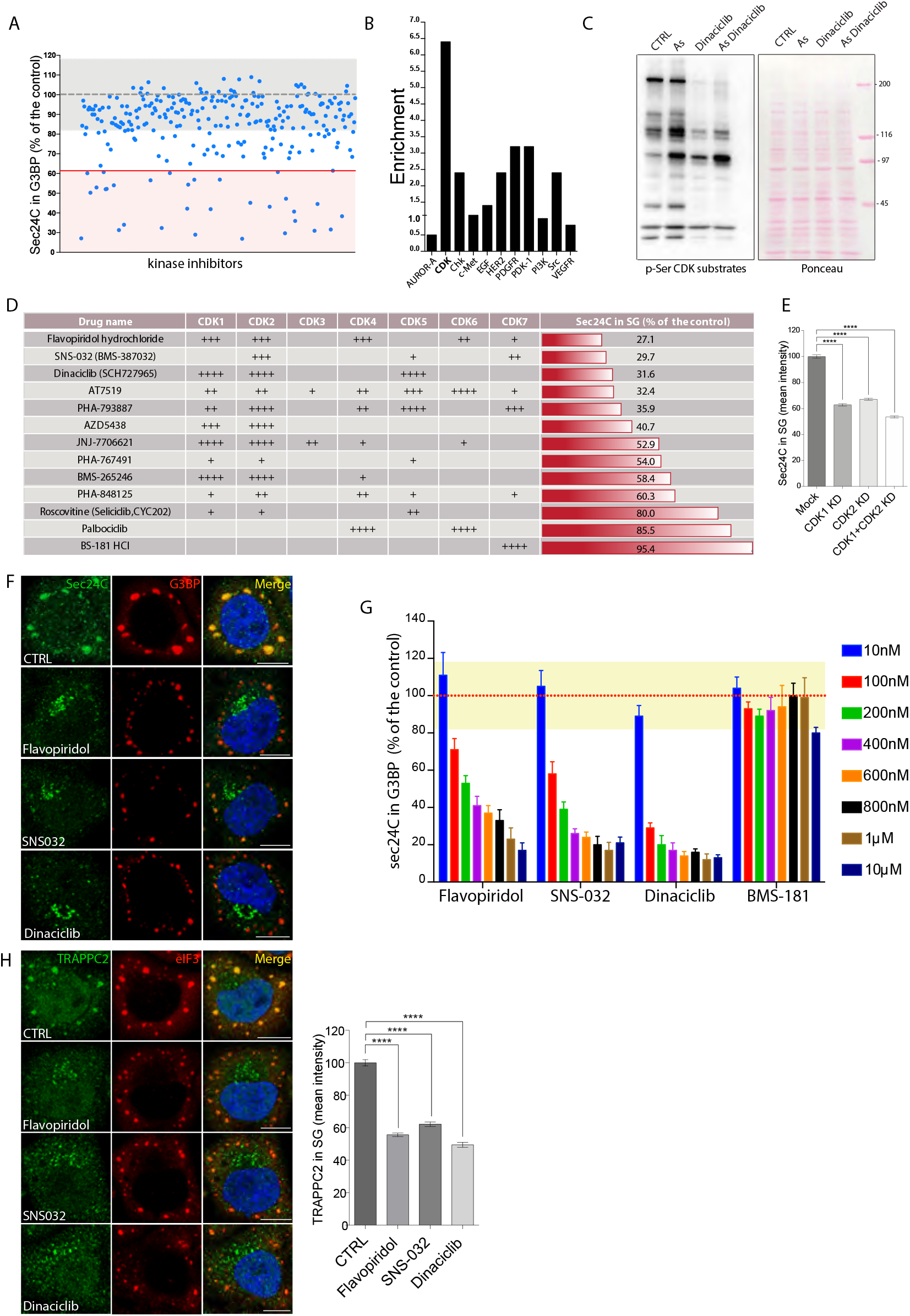
The association of TRAPP with SGs is under control of CDK1/2. **(A)** Cells were treated with 273 kinase inhibitors (10 μM) from the Sellekchem library and with SA, and Sec24C localization in G3BP puncta (% of total G3BP area) was calculated (see Materials and methods) and is reported as a percentage of the control (cells treated with SA alone). Gray dashed line, mean of control; gray box, control standard deviation (± 18.2%); red line, threshold of positive hits; red box: positive hits. **(B)** Enrichment Analysis of the positive hits; for each class of inhibitor, the enrichment (in percentage) of compounds falling into positive hits and the enrichment (in percentage) in the total number of compounds was calculated and expressed as a ratio. **(C)** Evaluation of CDK kinase activity upon SA treatment. Western blot analysis of phosphoSer-CDK substrates in control, SA (300μM, 30 min), Dinaciclib (10 μM, 180 min) and SA+Dinaciclib (150 min Dinaciclib and 30 minutes SA). CDKs are hyperactivated upon oxidative stress and this activation is partially prevented upon CDK inhibition. Left, Western blot, right: Ponceau was used as loading control. Image, one representative experiments out of 3 independent replicates. **(D)** Drug specificity, from high (++++) to low (+). for the different CDKs as described by Selleckchem, and Sec24C recruitment to SGs as a percentage of control. **(E)** Analysis of Sec24C recruitment to SGs in control (MOCK), CDK1-KD, CDK2-KD and CDK1+2-KD HeLa cells. Data are expressed as percentage of Sec24C (mean intensity) in SG puncta normalized for cytosolic signal as a percentage of control conditions. Quantification of one representative experiment. N>100, mean ± s.e.m. ****p<0.0001. **(F)** Representative images of HeLa cells treated with the best three hits (Flavopiridol, SNS-032, Dinaciclib; 1μM 150 min) and then exposed to SA (300 μM, 30 min), followed by immunostaining for Sec24C and G3BP. Scale bar, 10μm. **(G)** HeLa cells were treated with Flavopiridol, SNS-032, or Dinaciclib for 150 min at the indicated concentrations, subsequently treated with SA (300 μM, 30 min), and then immunostained for Sec24C and G3BP and imaged by OPERA. BS-181, a specific CDK7 inhibitor, was used as negative control. Sec24C localization to SGs is expressed as a percentage of the control (cells treated with SA alone). Dashed red line, mean of control; yellow box, standard deviation of control (± 9.8%). **(H)** Cells were treated as in (G) and immunostained for TRAPPC2 and eIF3. Scale bar, 10 μm. The graph shows the quantification of TRAPPC2 localization in SG puncta (mean intensity). Data are mean ± s.e.m. expressed as a percentage of TRAPPC2 signal in SGs after Flavopiridol, SNS-032 or Dinaciclib treatment compared to the control (cells treated with SA alone). N=3, three independent experiments, n=60-80 cells per experiment. ****p<0.0001. **Figure 4—figure supplement 1.** Evaluation of CDK1 and CDK2 knock down efficiency.

The CDK protein kinase family is composed of 21 members (Malumbres, 2014) and available inhibitors have limited selectivity (Asghar et al., 2015). To identify which specific CDK might be involved in Sec24C recruitment, we compared the potency of selected CDK inhibitors (Flavopiridol hydrochloride, SNS-032, Dinaciclib, AT7519, PHA-793887, ADZ5438, JNJ-7706621, PHA-767491, BMS-265246, PHA-848125, Roscovitine, Palbociclib, BS-181HCl) in the Sec24C recruitment assay with their ability to inhibit different CDK isoforms, as described by Selleckchem (https://www.selleckchem.com/CDK.html). This analysis revealed that CDK1 and 2 were those commonly targeted by the inhibitors that most potently affected Sec24C recruitment to SGs (Figure 4D), strongly suggesting that CDK1/2 are the most likely CDKs controlling COPII recruitment to SGs. This was confirmed by the observation that down-regulating the expression of CDK1 and CDK2 by siRNA inhibited COPII recruitment to SGs (Figure 4E, Figure 4— figure supplement 1). The three best hits (Flavopiridol, Dinaciclib, SNS032, Figure 4D,F) were assayed using a dose-response analysis and were found to be effective at concentrations as low as 100 nM (Figure 4G). As a negative control, the specific CDK7 inhibitor (BS-181HCl) was ineffective, even at the highest concentration tested (Figure 4G).

Since COPII recruitment to SGs is driven by TRAPPC2 recruitment (Figure 2F), we predicted that CDK1/2 inhibitors would also inhibit TRAPP recruitment. In line with our expectations, TRAPPC2 re-localization to SGs was reduced in cells treated with CDK1/2 inhibitors (Figure 4H).

Altogether, these results indicate that a signaling pathway involving CDK1/2 regulates the redistribution of TRAPP and COPII from ERES to SGs and prompted us to test the hypothesis that CDK inhibition might impair the recruitment of TRAPP/COPII to SGs by stabilizing their association with ER membranes. Indeed, we found that CDK inhibitors stabilize TRAPP and COPII on ERES membranes even in unstressed cells (Figure 5A). The analysis of the ERES associated pool using a permeabilization assay confirmed that the CDK1/2 inhibitors strongly stabilized COPII at the ERES, thus increasing the membrane associated pool and decreasing the cytosolic one (Figure 5B,C).

**Figure 5.**
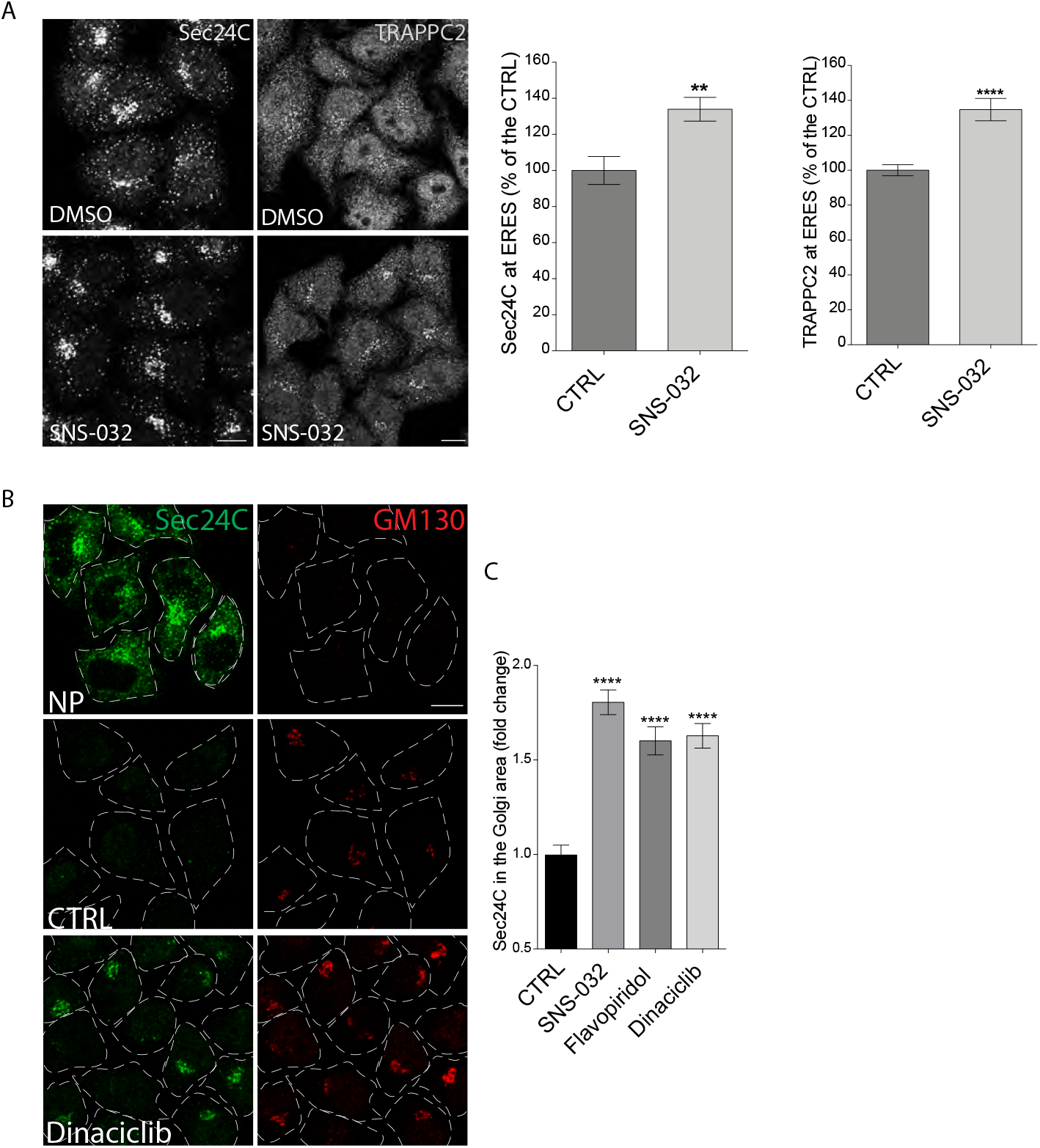
CDK1/2 modulate the COPII cycle at ERES. **(A)** Immunofluorescence images of Sec24C and TRAPPC2 in vehicle-treated and SNS-032-treated (1 μM, 3 h) cells. Scale bar, 10 μm. Graphs show the quantification of Sec24C and TRAPPC2 in the peri-Golgi area (mean intensity) normalized in SNS-032-treated cells relative to vehicle-treated cells (set as 100%). Mean ± s.e.m. of three independent experiments. **p<0.05, ****p<0.0001. **(B)** HeLa cells were treated with Dinaciclib (1 μM, 3 h) and permeabilized or not with digitonin as described in Materials and methods. A GM130 antibody was added to the buffer of living cells to monitor permeabilization efficiency. Upper panels, non-permeabilized (NP) control cells; middle panels, permeabilized control cells; lower panels, permeabilized Dinaciclib-treated cells. Scale bar, 10 μm. **(C)** Quantification of Sec24C membrane association after CDK inhibitor treatment. Digitonin-permeabilized cells treated with vehicle (CTRL) or the indicated CDK inhibitor (1 μM, 3 h) were immunostained for Sec24C. The mean intensity of Sec24C in the perinuclear area normalized for the cytosolic Sec24C signal in drug-treated cells is expressed as fold change compared to the control; n= 60-80; mean ± s.e.m. of three independent experiments ****p<0.0001. Scale bars, 10 μm.

Altogether, the above results suggest that at least one of the mechanisms underlying the regulation of TRAPP/COPII recruitment to SGs by CDK1/2 involves the ability of CDK1/2 to control the rate of COPII, and TRAPP cycling at the ERES.

### The recruitment of TRAPP and COPII to SGs occurs in actively proliferating cells

The observation that the recruitment of TRAPP and COPII to SGs is under control of CDK1/2 suggested that it could be linked to the cell proliferation. We explored this possibility by decreasing the cell proliferation rate in diverse and independent ways.

First, cells were subjected to prolonged nutrient starvation (HBSS for 8 hr) and then were exposed to SA for 30 min. Under these conditions, which inhibited cell proliferation, TRAPP/COPII association with SGs was reduced (Figure 6A,B). Second, HeLa cells were seeded at different confluency and then treated with SA for 30 min. Strikingly, the extent of confluency was inversely related to the extent of TRAPP/COPII association with SGs and to the degree of proliferation (Figure 6C).

**Figure 6.**
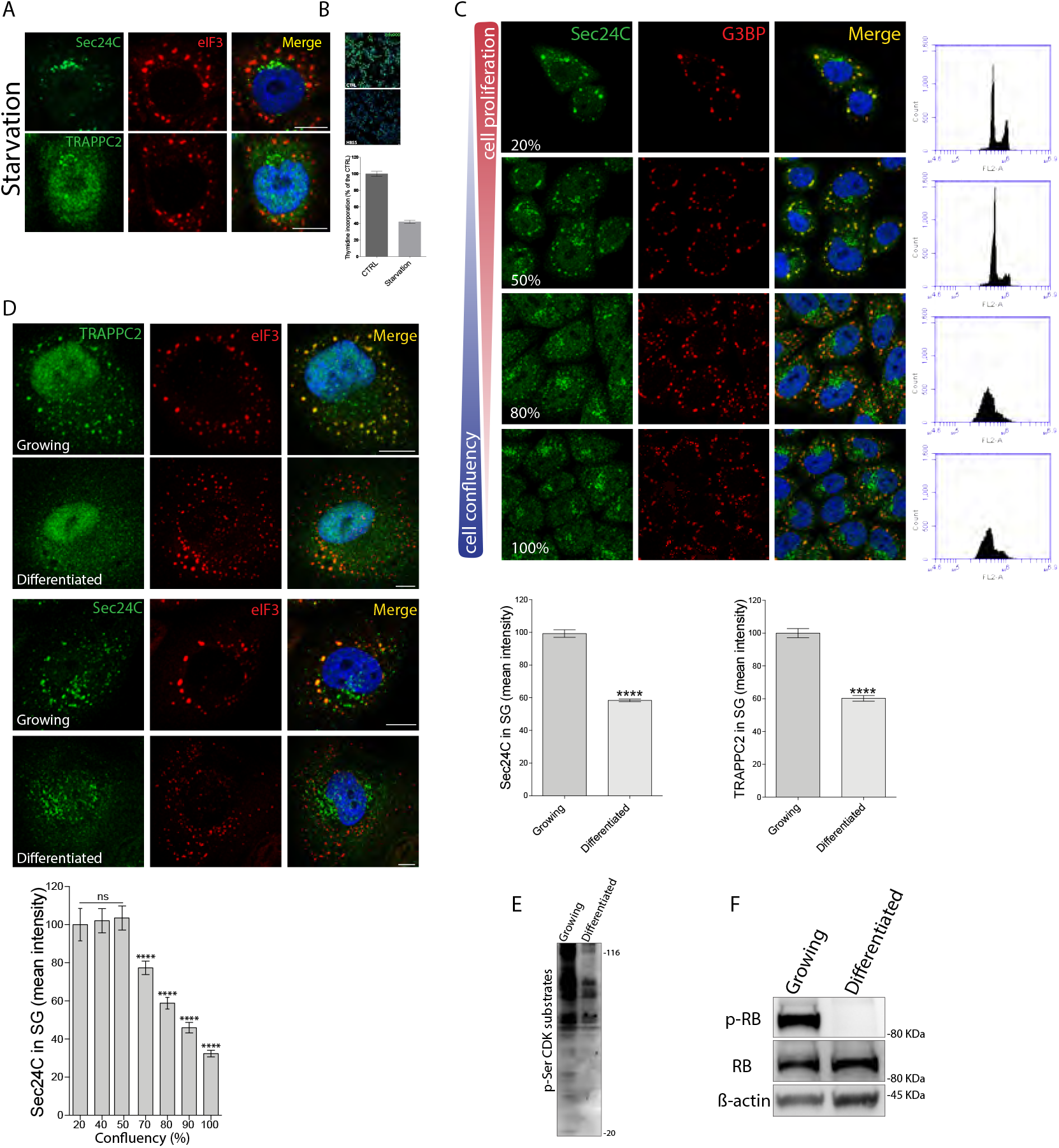
TRAPP and Sec23/24 migration to SGs depends on the proliferation state of cells. **(A)** HeLa cells were starved for 8 h with HBSS and subsequently exposed to SA (30 min, 300 μM), and immunostained for Sec24C or TRAPPC2 and eIF3. Scale bar 10 μm. **(B)** Analysis of the proliferation status of the cells after 8 h starvation in HBSS. Left, representative images of starved and nonstarved cells using EdU incorporation (see Materials and methods) and, right, quantification of EdU incorporation in starved cells as a percentage of incorporation in control fed cells. **(C)** HeLa cells were seeded at different confluency, treated with SA and stained for Sec24C and G3BP as an SG marker. Scale bar, 10 μm. Flow cytometry (FACS) analysis (right panels) was performed to evaluate the distribution of cell cycle phases in HeLa cell populations seeded at different confluency. The graph shows quantification of Sec24C mean intensity in SG puncta normalized for the cytosolic Sec24C at the indicated cell confluency. Mean ± s.e.m. of a representative experiment out of n=5 biological replicates. n= 50-80. ns, not significant; ****p<0.0001. **(D)** Growing and differentiated podocytes were treated with SA (300 μM, 30 min) and stained for TRAPPC2 or Sec24C and eIF3. Scale bar, 10μm. The graphs show quantification of TRAPPC2 and Sec24C (mean intensity) in SGs. Mean ± s.e.m. of three independents experiments ****p<0.0001. **(E)** CDK kinase activity in growing and differentiated podocytes. Western blot using a specific antibody recognizing phosphoSer-CDK substrates was used on total cell lysates. **(F)** Western blot analysis of phosphorylated retinoblastoma (p-RB) in growing (non-differentiated) versus differentiated podocytes. ß-actin was used as loading control. **Figure 6—figure supplement 1.** Differentiation of podocytes.

Third, we monitored the recruitment of TRAPP/COPII to SGs in podocyte cells expressing a temperature sensitive LTA-SV40 that actively proliferate at the permissive temperature of 33°C but arrest proliferation and differentiate at the restrictive temperature of 37°C (Saleem et al., 2002) (Figure 6—figure supplement 1). While TRAPP and COPII associate with SGs in undifferentiated podocytes in response to stress, this association was reduced in differentiated podocytes (Figure 6D). Of note, the proliferation to differentiation switch, as described (Saurus et al., 2016), is also accompanied by a decline in CDK activity (Figure 6E,F).

Altogether these data indicate that the association of COPII and TRAPP with SGs occurs only in proliferating cells and requires active CDK1/2 signaling.

### The TRAPP complex imposes SG size, composition, and function

The identification of TRAPP as a component of SGs led us to ask whether it could play any role in the assembly and function of SGs themselves. Indeed, we observed that depleting the entire TRAPP complex or only TRAPPC2 (through siRNA-mediated depletion) decreased the size of SGs (Figure 2F and Figure 7A). Microinjection of a TRAPPC3-blocking Ab (Yu et al., 2006; Venditti et al., 2012) had a similar effect (Figure 7B). Moreover, a smaller size of SGs was also observed upon treatment with CDK1/2 inhibitors, which prevents TRAPP translocation to SGs (Figure 4F, H). Under all of the above conditions the size of the SGs recall that of “immature SGs” (Wheeler et al., 2016).

**Figure 7.**
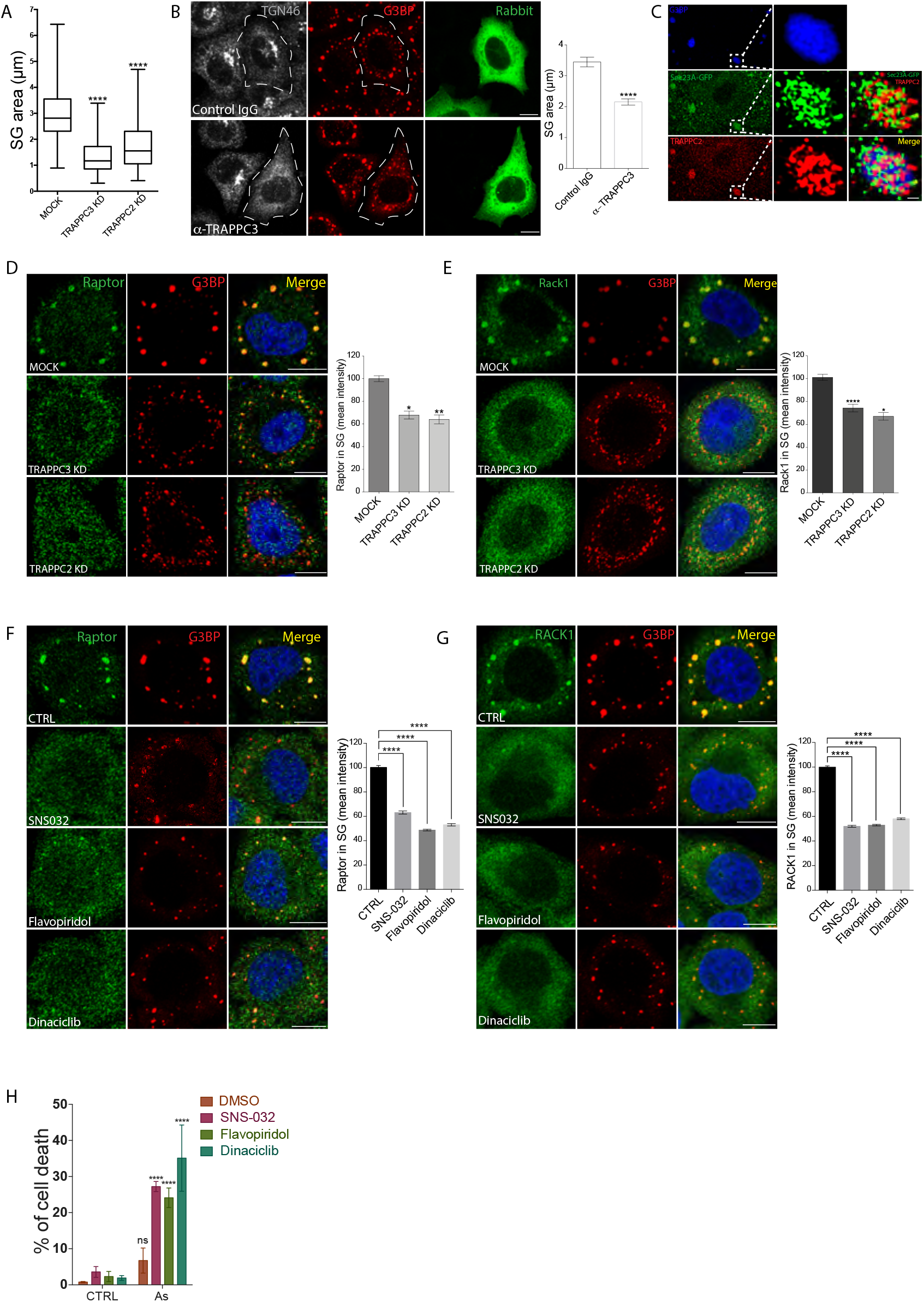
The TRAPP complex controls SG composition and function. **(A)** Box plot representing SG area (see Materials and methods) in mock, TRAPPC2- and TRAPPC3-depleted cells treated with SA. Distribution of values from three independent experiments. ****p<0.0001. **(B)** HeLa cells microinjected with control preimmune IgG or a TRAPPC3-specific antibody (right panels in green) were treated with SA. The TRAPPC3 Ab disrupts the Golgi (Yu et al., 2006), monitored using an anti-TGN46 Ab. Anti-G3BP was used to stain SGs. The graph shows quantification of the SG area. Mean ± s.e.m. three independent experiments; n>80. ****p<0.0001. Scale bar, 10 μm. **(C)** Structured Illumination Microscopy (SIM)-Super resolution (SR) images of endogenous TRAPPC2 and GFP-Sec23 localizing at SGs, stained for G3BP. Right, magnification of boxed area. Scale bar, 0.5 μm **(D,E)** Localization of Raptor **(D)** and RACK1 **(E)** in mock, TRAPPC3-KD and TRAPPC2-KD HeLa cells treated with SA. G3BP was used to stain SGs. Scale bar, 10 μm. Each graph shows the quantification (mean intensity) of the respective protein in SG spots expressed as a percentage of the mock. Mean ± S.D. three independent replicates. *p<0.02; **p<0.009 in **(D)**, *p<0.05; ****p<0.0001 in **(E). (F,G)** Localization of Raptor **(F)** and RACK1 **(G)** in untreated cells or cells pretreated with the indicated CDK inhibitor (1 μM, 150 min) and then with SA (300 μM, 30 min). G3BP was used to stain SGs. Scale bar, 10 μm. Graphs show quantification of the localization of the respective protein with SGs, expressed as a percentage of the control. Mean ± s.e.m. of one representative experiment out of three independent replicates, n=60-80. ****p<0.0001. **(H)** Analysis of cell death after overnight recovery of HeLa cells treated or untreated with CDKi (SNS-032; Flavopiridol or Dinaciclib, 1 μM, 150 min) and then treated with SA (500 μM, 3 h). Images were automatically acquired by OPERETTA microscope. Values indicate the percentage of the total number of nuclei (stained with DAPI) positive for BoBo-3 staining. Mean ± s.d. of one representative experiment out of three independent replicates. ns, not significant, ****p<0.0001. **Figure 7—figure supplement 1.** TRAPP depletion does not affect protein translation inhibition caused by SA treatment.

In fact, the processes that lead to SG nucleation, i.e. shut-down of protein synthesis or phosphorylation of eIF2alpha induced by oxidative stress, were not affected by TRAPP depletion (Figure 7—figure supplement 1A,B). Instead, the successive increase in size of the nucleated SGs (i.e. SG maturation) was impaired in TRAPP-depleted cells, consistent with our observation that TRAPP is recruited after the initial formation of SGs (Figure 1D) and occupies an outer position in the SGs (Figure 7C). A feature of SG maturation is the acquisition of an array of signaling molecules (which can be either hyperactivated or deactivated) to drive the pro-survival program (Franzmann and Alberti, 2019). This is the case of Raptor, that controls the energy sensor mTORC1 and of the kinase RACK1 (Arimoto et al., 2008; Thedieck et al., 2013; Wippich et al., 2013). To evaluate whether the absence of TRAPP could affect SG maturation and/or signaling function by interfering with the recruitment of these proteins, we checked the localization of Raptor (Figure 7D) and RACK1 (Figure 7E, Figure 7—figure supplement 1C) in TRAPP- and or TRAPPC2-depleted cells subjected to oxidative stress. We observed that the amount of these signaling proteins sequestered to SGs was strongly reduced by TRAPP depletion, thus indicating that the TRAPP complex is necessary to generate “mature” SGs that can accomplish a fully functional stress response. Localization of RACK1 and Raptor was also evaluated in a context of CDK inhibition. Again, treatment with CDK inhibitors strongly reduced Raptor (Figure 7F) and RACK1 (Figure 7G) sequestration to SG, further supporting that the TRAPP complex is needed for SG maturation. Notably, the distribution of Raptor in response to oxidative stress was also studied by transiently expressing a cMyc-tagged version of the protein in control, in TRAPP-KD cells and in cells treated with CDK inhibitors. While myc-Raptor readily relocated to SGs in control cells exposed to stress, it failed to associate with SGs under conditions that prevented TRAPP association with SGs (i.e. TRAPP depletion and CDK inhibition; data not shown).

SGs are part of a pro-survival program that aims to save energy to cope with stress insults. Interfering with this program by impairing SG formation sensitizes cells to apoptosis (Takahashi et al., 2013). We thus assayed stress resistance in cells with an impaired capability to recruit TRAPP to SGs, i.e. upon CDK inhibition. As shown in Figure 7H, CDK inhibition significantly sensitized cells to cell death, suggesting that the presence of TRAPP at SGs is required for their anti-apoptotic role and highlighting the TRAPP complex as a novel player in the cell response to stress.

### The recruitment of TRAPP and COPII to SGs impairs ER export of newly synthesized cargoes and induces the fragmentation of the Golgi complex

We finally investigated the functional relevance of the recruitment of TRAPP/COPII to SGs by analyzing its impact on the function and structure of the secretory pathway.

We first analysed the ER export of procollagen type I (PC-I), which is known to be regulated by COPII and TRAPP (Gorur et al., 2017; Venditti et al., 2012). PC-I transport can be synchronized by temperature, being unfolded and blocked in the ER at 40°C while properly folded and assembled and able to be exported from the ER, transported to the Golgi complex (GC) and secreted at 32°C (Mironov et al., 2001, and Methods). Human fibroblasts were incubated at 40°C, left untreated or treated with SA, which induces the recruitment of Sec24C to SGs (Figure 8A). PC-I was blocked in the ER under both conditions (Figure 8A). Monitoring of PC-I transport after the shift to 32°C showed that while PCI reaches the GC in untreated cells, it remains blocked in the ER in SA-treated cells (Figure 8A).

**Figure 8.**
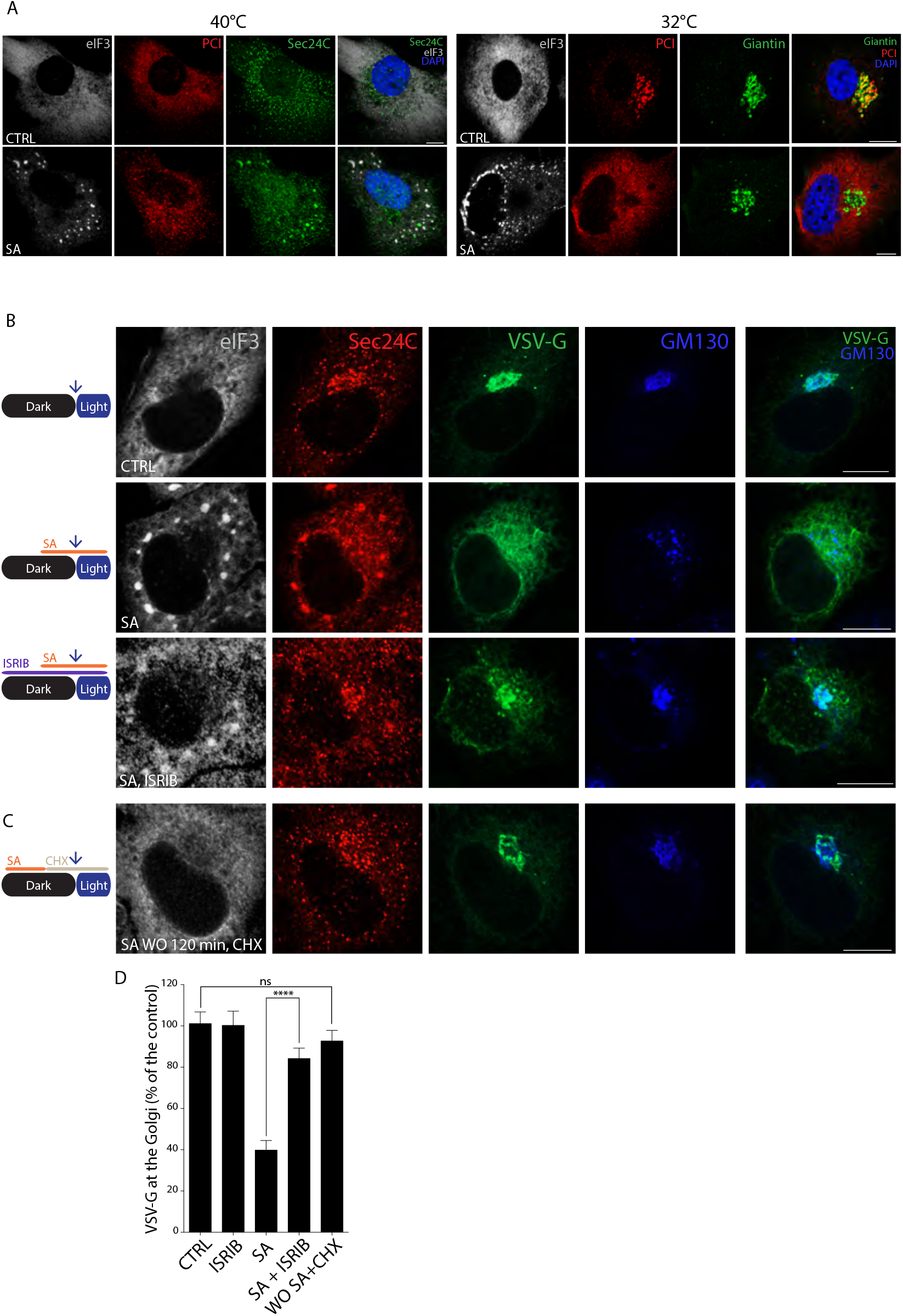
The sequestering of TRAPP and COPII in SGs slows down ER export. **(A)** Left, control or SA-treated (500μM, 120 min) human fibroblasts (HFs) were incubated at 40°C for 180 min and stained for eIF3, PCI, and Sec24C. Right, the cells were imaged 10 min after shifting the temperature from 40°C to 32°C and stained for eIF3, PCI, and Giantin (to label the Golgi). Scale bar, 10 μm. **(B)** HeLa cells expressing VSV-G-UVR8 mEOS were left untreated or treated with SA alone or in combination with ISRIB (1 μM) for the indicated times and then pulsed with blue light. Images were taken 10 minutes after the UV pulse and processed for staining. **(C)** Cells were treated with SA for 30 min, the SA was washed out (WO), and cells were left to recover for 180 min in the presence of cycloheximide, followed by a blue light pulse and processing for staining as described in (B). **(D)** Quantification of VSV-G (mean intensity) in the Golgi area to the total VSV-G per cell under the conditions described in (B,C). Data are expressed as percentage of the control. Mean ± s.e.m. of three independent experiment. ****p<0.0001; ns, not significant. **Figure 8—figure supplement 1.** ISRIB delays SG nucleation and inhibits Sec24C relocalization to SGs.

We then investigated to what extent the inhibition of cargo export from the ER induced by oxidative stress is dependent on COPII/TRAPP sequestering onto SGs. To this end we compared the ER export of neosynthesised cargo in control cells and in SA-treated cells either under conditions permissive for the formation of SGs (and of the recruitment of COPII/TRAPP to SG) or under conditions that prevent SG formation. For the latter, we took advantage of the use of ISRIB, a compound that antagonises the effect of eIF2α phosphorylation on translation and impairs SG assembly (Sidrauski et al., 2015). In preliminary experiments, we checked the ability of ISRIB to impair SG formation under our experimental conditions and found that ISRIB completely abrogates SG assembly after 15 min SA treatment, delaying their appearance at later time points of treatment and thus significantly also impairing COPII sequestering to SGs (Figure 8—figure supplement 1). To monitor the synchronized export of neosynthesised cargo we followed a reporter cargo, VSVG-UVR8, that exits the ER in a UV light-inducible manner (Chen et al., 2013). After UV light induction, VSVG-UVR8 exited the ER and reached the Golgi in control cells, while its ER export was severely impaired in cells treated with SA under conditions permissive for SG formation where COPII was massively recruited to SGs (Figure 8B, D). Interestingly, the inhibition of ER export of VSVG-UVR8 was largely prevented when the cells were exposed to similar oxidative stress, but under conditions that impaired SG formation (i.e. when cells were pretreated with ISRIB (Figure 8B, D)).

We next tested whether COPII and TRAPP maintain their functionality once released from SGs upon stress removal (Figure 1—figure supplement 3). Cells were treated with SA for 30 min, the SA was washed out, and cells were left to recover for 180 min in the presence of cycloheximide to prevent *de novo* synthesis of TRAPP and COPII components. Under these conditions SGs were resolved, COPII returned to its native location (ERES/cytosol) and cells completely recovered their capability to transport cargo to the Golgi apparatus (Figure 8C,D). These data indicate that sequestration of COPII/TRAPP onto SGs halts ER-to-Golgi trafficking while removal of the stress releases COPII/TRAPP and allows trafficking to resume.

COPII and TRAPP not only control ER export but are also needed to maintain the organization of the GC. In particular, the TRAPP complex acts as GEF for Rab1, a GTPase with a key role in the organization and function of the GC (Tisdale et al., 1992; Wilson et al., 1994). We monitored Golgi complex morphology in response to SA by following the distribution of the early Golgi marker GM130. Strikingly, the GC starts to fragment after 30 min and completely redistributes throughout the cytoplasm after 60 min of SA treatment (Figure 9A). This time window overlaps with the progressive recruitment of the TRAPP complex onto SGs (Figure 1D). To assess whether the Golgi fragmentation induced by oxidative stress was due to sequestration of TRAPP/COPII onto SGs, we analyzed the morphology of the GC in cells exposed to oxidative stress under conditions that prevent COPII/TRAPP recruitment to SGs (i.e. treatment with CDK inhibitors, highly confluent cells) or that impair SG formation (i.e. treatment with ISRIB). We found that CDK inhibitors prevented the fragmentation of the GC in cells exposed to oxidative stress (Figure 9B) and that oxidative stress had no impact on the organization of the Golgi complex in highly confluent cells (Figure 9C). Finally, ISRIB was effective in preventing the fragmentation of the Golgi complex induced by SA (Figure 9D).

**Figure 9.**
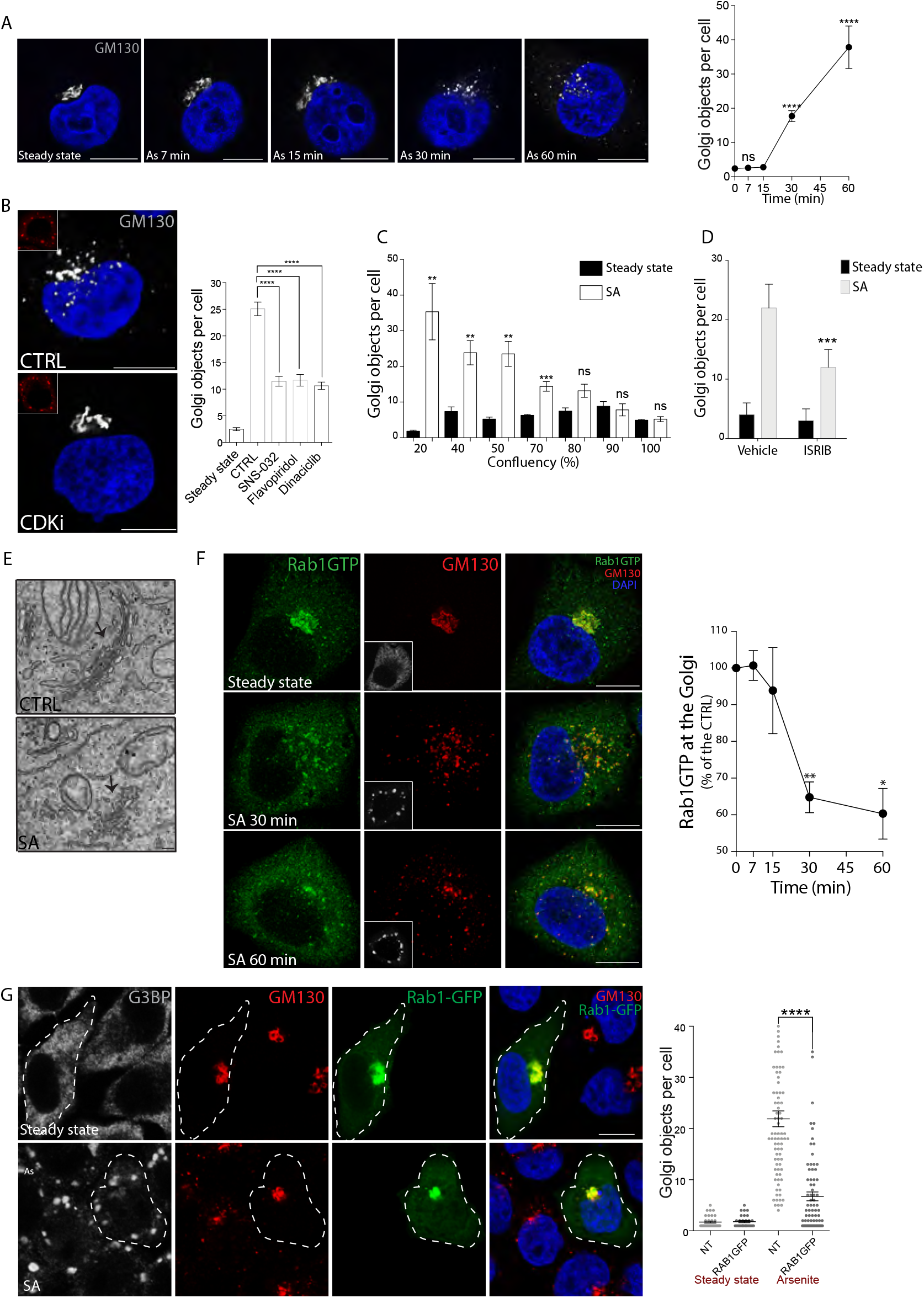
The sequestering of TRAPP in SGs induces the fragmentation of the GC in a Rab1-dependent manner. **(A)** AiryScan images of HeLa cells treated with SA for the indicated times and stained with GM130 (gray). Scale bar, 10 μm. The graph shows quantification of Golgi particles in SA-treated cells. Mean ± s.e.m. of three independent experiments. n=90-100. ****p<0.0001; ns, not significant. **(B)** Control cells and cells treated with Dinaciclib (CDKi) were treated with SA and stained for GM130 (in gray). Insets, eIF3. Blue, nuclear DAPI staining. The graph shows quantification of Golgi particles in HeLa cells untreated or treated with SNS-032, Flavopiridol, or Dinaciclib for 150 min and treated with SA (300 μM, 30 min). Mean ± s.e.m. of three independent experiment. ****p<0.0001. **(C)** Quantification of Golgi particles in cells plated at different percentage of confluency. Mean ± s.e.m. of three independent experiment. ** p<0.001; *** p<0.0002; ns: not significant. **(D)** Quantification of Golgi objects in cells pre-treated with ISRIB (1 μM) for 3 h and exposed to SA (30 min) in the presence of ISRIB. *** p<0.002. **(E)** Electron microscopy images of a cell exposed to SA (300 μM, 30 min). GC: Golgi complex. Scale bar 200 nm. **(F)** AiryScan images of HeLa cells at steady state or exposed to SA for the indicated times. A Rab1-GTP specific antibody was used to monitor the pool of active Rab1 and GM130 to stain the Golgi complex. Insets, eIF3. The graph shows quantification of Rab1-GTP at the Golgi complex expressed as a percentage of the control. Mean ± s.e.m. of three independent experiments. *p<0.011, **p<0.002. **(G)** HeLa cells transfected or not with WT GFP-Rab1B were left untreated or treated with SA and stained for GM130 and G3BP to monitor SGs. Dashed white line, WT GFP-Rab1B transfected cells. The graph shows quantification of Golgi particles in the different conditions. NT: non-transfected. Mean ± s.e.m. of three independent experiment. ****p<0.0001; n=30-50. **Figure 9—figure supplement 1.** TRAPPC2 migrates to SGs in Rab1 overexpressing cells.

These results indicated a causal link between TRAPP/COPII sequestration to SGs and Golgi fragmentation. To gain insight into the nature of the fragmentation of the Golgi complex, we performed ultrastructural analysis that revealed the presence of residual swollen cisternae with the loss of stacked structures and extensive vesiculation in SA-treated cells (Figure 9E), a situation that is reminiscent of the effect of Rab1 inactivation (Wilson et al., 1994). We hypothesized that the dismantling of the Golgi complex could have been due to impaired Rab1 activation as a consequence of TRAPP (the Rab1 GEF) sequestering to SGs. We monitored Rab1 activity by using an antibody that specifically recognizes the GTP-bound Rab1 and found, indeed, that the fraction of active Rab1 (i.e. the GTP-bound form) is progressively reduced in cells exposed to SA (Figure 9F). Overexpression of wt Rab1 reduced fragmentation/vesiculation of the GC induced by oxidative stress without affecting the capacity of TRAPP to migrate to SGs (Figure 9G, Figure 9—figure supplement 1), establishing a causal a link between Rab1 inactivation and Golgi fragmentation. It is worth mentioning that a further contribution to the disorganization of the Golgi complex upon oxidative stress may derive from the delocalization of the TGN-located PARP12 to SGs, which, however, affects mainly late Golgi compartments (Catara et al., 2017).

## Discussion

We described a branch of the integrated stress response that is dependent on the assembly of SGs and that impinges on the early secretory pathway. A key player of this branch is TRAPP, a multimolecular complex that intervenes in multiple membrane trafficking steps.

We showed that TRAPP is massively recruited to SGs in response to different acute stress stimuli and in turn recruits components of the inner layer of the COPII complex Sec23-Sec24, the protein complex that drives the export of newly synthesized proteins from the ER, to SGs. The functional consequences of the recruitment of TRAPP/COPII to SGs are double-edged since on the one hand they impact on the function and organization of the secretory pathway while on the other they impact on the composition and function of the SGs (Figure 10).

**Figure 10.**
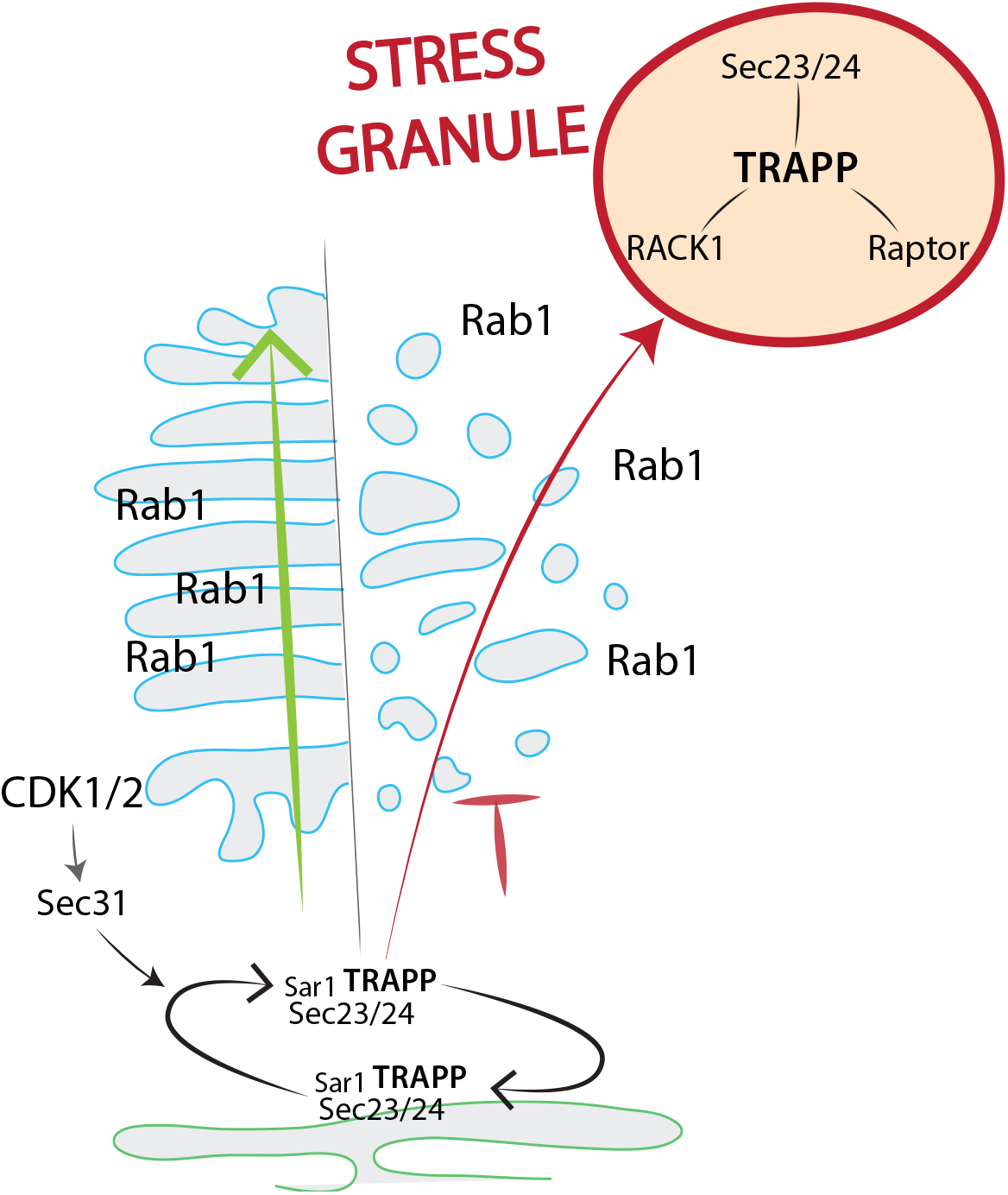
Proposed model for secretion arrest mediated by TRAPP upon SG assembly in actively proliferating cells. Upon stress, the TRAPP complex relocalizes to SGs and drives the sequestering of Sec23/24, RACK1 and Raptor to SGs. This phenomenon leads to a block in ER-export and contributes to SG maturation.

We have shown that the sequestering to SGs of these two complexes, one, COPII, with a pivotal role in ER export and another, TRAPP, acting as a GEF for Rab1, a master GTPase orchestrating the organization of the Golgi complex, induces a slow-down of the ER-to-Golgi transport of newly synthesized proteins and the disorganization of the structure of the Golgi complex. This halt in secretory activity, an energetically costly process, can be seen as a way to limit energy expenditure upon acute stress but also to prevent the ER export of newly synthesized proteins that might have been damaged/misfolded by the acute stress before an adequate UPR is installed and becomes operative. Importantly, these changes are completely reversible since ER export resumes and proper Golgi morphology is restored upon stress removal and relocation of the two complexes to their natural sites.

An intriguing feature of this branch of the integrated stress response that involves the secretory pathway is its strict dependence on the proliferation status of the cells. Indeed, the sequestration of TRAPP and COPII to SGs occurs only when the acute stress affects actively proliferating cells. Cells with a slow/halted proliferation rate still form SGs in response to acute stress stimuli, but do not delocalize TRAPP/COPII from their steady state sites, i.e., the ERES, to SGs. We have used multiple strategies to slow down (or halt) proliferation: nutrient deprivation, a high cell density, or a switch from a proliferation to a differentiation program. Under all circumstances, we observed a very tight correlation between the extent of recruitment of TRAPP/COPII to SGs and the proliferation rate, indicating that acquisition of components of the secretory machinery does not occur by default as a consequence of the assembly of the SGs but is an active process that is subject to specific regulation.

We found that this regulation involves the activity of CDK1&2, since CDK1&2 inhibitors and/or CDK1&2 downregulation prevented the relocation of TRAPP/COPII to SGs by stabilizing their association with the ERES. Indeed, the function of the ERES and some of the components of the COPII coat are known to be controlled by CDKs. A member of the CDK family, PCTAIRE, is recruited to ERES via its interaction with Sec23 and controls ER export (Palmer et al., 2005), while Sec31, a component and a regulator of Sar1/COPII cycle, can be phosphorylated by CDKs (Holt et al., 2009; Hu et al., 2016). Further, the phosphorylation of Sec31 in response to growth factor and mitogenic signals has also been reported (Dephoure et al. 2008; Olsen et al., 2006). We have shown here that, consistent with the described role of Sec31 as a co-GAP for Sar1, Sec31 function is required to control the COPII cycle at the ERES, since Sec31 depletion induces a tighter association of Sec23-24 with ERES and prevents their translocation to SGs.

In fact, our results indicate that the extent of translocation of COPII/TRAPP to SGs is directly proportional to their cycling rate at the ERES membranes. We have shown that this rate is diminished by CDK inhibitors, but also, importantly, in starved cells as compared to actively proliferating cells. These observations are in line with previous reports showing that the cycling rate of ERES components is under control of growth factors and growth factor-dependent signaling (Farhan et al., 2010; Tillmann et al., 2015). Thus, the activity of the ERES, as measured by their number and by the rate of cycling of COPII and other ERES components, is stimulated under conditions that demand maximal output efficiency from the ER (i.e. during cell proliferation) as they involve cell size increase and organelle expansion. If cells experience an acute stress under these conditions of active growth, then the fast cycling COPII components are susceptible to being temporarily sequestered to the SGs to instantly slow down ER output and reduce energy consumption. By contrast, conditions that are not permissive for cell proliferation, such as nutrient starvation, stabilize the association of COPII/TRAPP with ERES membranes and prevent their translocation to SGs. Indeed, a profound remodeling of ERES is known to occur during nutrient starvation and coincides with the rerouting of COPII and TRAPP from their role in mediating the ER export of newly synthesized proteins to their role in autophagosome formation (Ge et al., 2017; Imai et al., 2016; Kim et al., 2016; Lamb et al., 2016; Ramírez-Peinado et al., 2017; van Leeuwen et al., 2018). The kinetics of the COPII cycling at the sites of phagosome formation in the ER have not been specifically explored. It is tempting to speculate that it might be slower than that at the “conventional” ERES under feeding conditions and that this more stable association of COPII with the ER during autophagosome formation may in turn prevent its sequestration to SGs upon stress exposure. Preventing the sequestration of TRAPP/COPII in starved cells exposed to stress would preserve the function of COPII and TRAPP in the autophagy process, a process that leads to nutrient recycling and energy production, two desirable events in cells exposed to stress.

Finally, we have shown that TRAPP is required for the proper maturation of SGs. Impeding the recruitment of TRAPP to SGs or depleting TRAPP does not impair the formation of SGs *per se* but hampers their maturation, as evaluated by their size (smaller SGs in the absence of TRAPP) and composition. In particular, key signaling components, such as RACK1 and Raptor, are no longer recruited to SGs in TRAPP-depleted cells. The mechanisms underlying the TRAPP-dependent maturation of SGs remain to be identified. Two possible non-exclusive scenarios are worthy of consideration, the engagement of specific protein-protein interactions between TRAPP components and signaling molecules or a central role of TRAPP, originally identified as a tethering complex, in mediating tethering/coalescence of smaller SGs into larger ones, which might be the only ones competent for recruiting signaling molecules. Whatever the underlying mechanisms, the impaired recruitment to SGs of RACK1 and Raptor in TRAPP-depleted cells leads to lower resistance to stress and higher tendency to undergo apoptosis, as the association of these signaling elements with SGs exerts an anti-apoptotic role (Arimoto et al., 2008; Thedieck et al., 2013; Wippich et al., 2013). Our finding that TRAPP, and in particular TRAPPC2, acts as a mediator of the secretory arrest in response to stress and as a driver of SG maturation elicits the question as to whether and how these unsuspected properties of TRAPPC2 may be relevant for the pathogenesis of SEDT, which is caused by mutations in TRAPPC2 (Gedeon et al., 1999). This, as well as the more general question on the physiological relevance of assembling SGs in response to stress (Protter and Parker, 2016), remain outstanding questions for further investigation. For now, we can only speculatively propose that, given the role of TRAPPC2 in mediating secretory arrest and SG maturation in actively proliferating cells, the main target cells in SEDT are likely to be proliferative chondrocytes. Derived from chondrocyte progenitor cells, rapidly proliferating chondrocytes are key components of the epiphyseal growth plate and undergo terminal differentiation into postmitotic hypertrophic chondrocytes, which eventually transdifferentiate into osteoblasts or undergo apoptosis, being replaced by osteoblasts and osteoclasts. It is established that the rate of proliferation, together with the height of columnar hypertrophic chondrocytes, are major determinant of growth plate function and thus of bone length (Ballock and O’Keefe, 2003). We hypothesize that TRAPPC2-defective proliferative chondrocytes are less resistant to repetitive stresses, including mechanical and oxidative stresses (Henrotin et al., 2003; Zuscik et al., 2008), and are thus more prone to undergo apoptosis. The accelerated loss of proliferative chondrocytes would thus lead to a premature growth plate senescence that could manifest as progressive delayed growth.

## Supplementary Materials

### Materials and methods

#### Reagents and antibodies

Primary antibodies used in this study were: mouse monoclonal antibody anti-G3BP (BD Transduction Laboratories cat. no. 611126), rabbit polyclonal antibody anti-G3BP (Bethil, cat. no. A302-033), mouse monoclonal antibody anti-TRAPPC5 (Abnova cat. no. H00126003-A01), goat polyclonal antibody anti-eIF3 (Santa Cruz cat. no. sc-16377), rabbit polyclonal antibody anti-Sec24C (Sigma-Aldrich cat. no. HPA040196), mouse monoclonal anti-TIAR (BD Transduction Laboratories cat. no. 610352), mouse monoclonal TRAPPC12 (Abcam cat. no. 88751), mouse monoclonal TRAPPC10 (Sanza Cruz cat. no. SC-101259) mouse monoclonal antiflag (Sigma-Aldrich cat. no. F1804), rabbit polyclonal antibody anti-phosphor-Ser CDK substrates motif (Cell signaling cat. no. 9477), mouse monoclonal anti-GM130 (BD Transduction Laboratories cat. no. 610823), human monoclonal antibody anti-Rab1-GTP (Adipogen cat. no. AG-27B-0006), mouse monoclonal anti-RACK1 (Sanza Cruz cat. no. 17754), rabbit polyclonal anti-Raptor (Cell signaling cat.no. 2280), rabbit polyclonal Sec16A (Bethil, cat.no. A300-648A), sheep polyclonal anti-TGN46 (Serotec cat. no. AHP500), rabbit polyclonal antibody anti-Sec31A (Sigma-Aldrich cat. no. HPA005457), mouse monoclonal antibody AP-II (Pierce, cat. no. MA1-064), mouse monoclonal *γ*-adaptin (Transduction cat. no. 618305), mouse monoclonal antibody anti-Retinoblastoma (Cell Signaling cat. no. 9309), rabbit polyclonal antibody anti-phospho Retinoblastoma (Cell Signaling cat. no. 9516), rabbit polyclonal antibody anti-eiF2α (Cell Signaling cat. no. 9722), rabbit polyclonal antibody anti-phospho-eiF2α (ser51) (Cell Signaling cat. no. 3597), mouse monoclonal antibody anti-puromycin (Millipore cat.no. MABE343), rabbit polyclonal anti-ß-actin (Sigma-Aldrich cat. no. A2066), mouse monoclonal antibody anti-GAPDH (Santa Cruz cat. no. sc-32233). Guinea pig polyclonal antibody anti-Synaptopodin (Acris cat. no. AP33487SU-N). Polyclonal antibodies against Arfaptin, p115, G-97, COPI, TRAPPC2 and TRAPPC3 were obtained in our labs. Media, serum and reagents for tissue culture were purchased from Thermo Fisher Scientific.

Sodium arsenite (cat. no. S7400), Puromycin (cat.no. A1113802) and cycloheximide (cat.no. C7698) were purchased from Sigma-Aldrich, Flavopirodol hydrochloride (cat. no. S2679), SNS-032 (BMS-387032) (cat. no. S1145) and Dinaciclib (SCH727965) (cat. no. S2768) were purchased from Sellekchem.

#### Plasmid construction

TRAPPC3-GFP, Sec23A-GFP, Sar1GTP-GFP, Rab1-GFP, OCRL-GFP, Rab5-GFP and Rab7-GFP were obtained as previously described (De Leo et al., 2016; Venditti et al., 2012; Vicinanza et al., 2011). Construct encoding GFP-ARF1 has been described (Dubois et al., 2005).

#### Cell culture, transfection, and RNA interference

HeLa cells were grown in high glucose (4,500 mg/L) DMEM supplemented with 10% FCS. U2OS were grown in McCoy’s supplemented with 20% FCS. Human fibroblasts were grown in DMEM and M199 (1:4) supplemented with 10% FCS. Podocytes were grown in RPMI-1640 supplemented with 10% Insulin-transferrin-Selenium (ITS) and 10% FCS.

For transfection of DNA plasmids, HeLa cells were transfected using either TransIT-LT1 (Mirus Bio LLC, for BioID2 experiment) or JetPEI (Polyplus, for immunofluorescence analysis) as transfection reagents, and the expression was maintained for 16 hr before processing. Microinjection of TRAPPC3 antibody was performed as described (Venditti et al., 2012).

siRNA sequences used in this study are listed in table S2. HeLa cells were treated for 72 hr with Oligofectamine (Life Technologies) for direct transfection. Knock-down efficiency was evaluated by western blot for TRAPPC2, TRAPPC3 and Sec31 (figure 2-Supplementary 3C and figure 3-Supplementary 1) and by q-PCR for Sec16, Sec23 and sec24 isoforms, CDK1 and CDK2 (figure 2-Supplementary 2; figure 2-Supplementary 3A,B; figure 4 Supplementary 1).

#### Generation of TRAPPC2 KO cell line

SEDL-KO HeLa cells were generated by double infection with lentiviral particles (LentiArray Particles, Thermo Fischer Scientific), one lentiviral containing a cassette for expression of a gRNA sequence to target both TRAPPC2 (chrX) and TRAPPC2B (chr19) “GTTCAACGAGTGGTTTGTGT” and the other to express Cas9. The cells were coinfected at 70% confluency with a MOI of 1:10 gRNA:CAS9 in the presence of 5ug/ml of polybrene (Sigma). After 48h the cells were incubated in medium containing 1μg/ml of puromycin (Calbiochem) and 0.5 μg/ml blasticidin (Invivogen). After a further 5 days, single-cell sorting in 96-well plates was performed using FACSAria III from Becton Dickinson (BD Biocsciences, San Jose, USA). 2-3 week later, single clones were detached and analyzed by PCR and subsequent sequencing. Positive clones were further verified by SDS-PAGE and immunofluorescence analysis.

#### Stress stimuli

Mammalian cells were treated with 300 μM SA for 30 min (unless otherwise stated) in DMEM 10% FBS supplemented with 20mM Hepes. Temperature was maintained at 37°C for the time of treatment in water bath. Heat treatment (44°C) was for 45 min in water bath.

#### Immunofluorescence microscopy

HeLa, human fibroblast, U2OS and rat chondrosarcoma cells were grown on coverslips and fixed with 4% PFA for 10 min, washed three times with PBS, blocked and permeabilized for 5 min in 0.1% Triton in PBS. Samples were washed three times and blocked with blocking solution (0.05% saponin, 0.5% BSA, 50 mM NH4Cl in PBS) and incubated with primary antibodies diluted in blocking solution for 1 hr at RT. Coverslips were washed with PBS and incubated with fluorochrome-conjugated secondary antibodies (Alexa-Fluor-488, Alexa-Fluor-568 and Alexa-Fluor-633) for 1 hr at RT. Fixed cells were mounted in Mowiol and imaged with Plan-Apochromat 63x/1.4 oil objective on a Zeiss LSM800 confocal system equipped with an ESID detector and controlled by a Zen blue software. Fluorescence images presented are representative of cells imaged in at least three independent experiments and were processed with Fiji (ImageJ; National Institutes of Health) software. Sec24C and TRAPPC2 quantification and Golgi particle analysis are described in the corresponding figure legends.

Differences among groups were performed using the unpaired Student’s *t* test calculated with the GraphPad Prism software. All data are reported as mean ± s.e.m. as indicated in the figure legends. For Structured Illumination Microscopy (SIM, Figure 7D, figure7-supplementary 1C), cells were imaged with a Plan-Apochromat 63x/1.4 oil objective on a Zeiss LSM880 confocal system. Image stacks of 0.88 μm thickness were acquired with 0.126 μm z-steps and 15 images (three angles and five phases per angle) per z-section. The field of view (FOV) was ~ 75 x 75 μm at 0.065 μm/pixel, and a SIM grating of 34 μm was used. Image stacks were processed and reconstructed with Zen software, using the SIM tool in the Processing palette.

#### Gel filtration

The lysate used for gel filtration was prepared by cell cracker homogenizing 1.8X10^8^ cells in 2 ml of buffer 20 mM HEPES, pH 7.2, 150 mM NaCl, 2 mM EDTA, 1 mM DTT and Protease inhibitor (Roche). The sample was centrifuged at 100,000xg for 1 h. 10 mg of protein was concentrated in 350μL and loaded onto a Superose6 gel filtration column (GE) and 400 μL of each fraction was collected. 50μL of fractions were processed for SDS-PAGE analysis and proteins were detected by Western blot using specific antibodies as described in Figure 1E.

#### Yeast Methods

The centromeric plasmid pUG23-Bet3-GFP (His selection) was described previously (Mahfouz et al., 2012). A Pab1-mRFP expressing plasmid was constructed by PCR amplification of the Pab1 gene plus 419 bp of the promoter region from yeast genomic DNA and cloning into the vector backbone of the centromeric plasmid pUG35 (Ura selection), removing the Met25 promoter and replacing GFP with mRFP, using standard cloning techniques. Yeast cells (BY4743) were transformed with both plasmids and grown in -Ura-His media with 2% Dextrose at 30°C to early log phase. Stress granules were induced by incubating cells at 46°C for 10 min (Riback et al., 2017) and observed immediately on an LSM800 microscope.

#### Immunoprecipitation

For the studies of TRAPPC2 interactors, HeLa cells were transfected with the TRAPPC2-3XFLAG and 3XFLAG constructs. 16 hr post-transfection, cells were lysed with lysis buffer (50 mM Tris-HCl pH 7.4, 100 mM NaCl, 1 mM EDTA, 0.5% Lauryl Maltoside, protease and phosphatase inhibitors).

Cell extracts from control transfected HeLa cells or cells expressing TRAPPC2-3XFLAG were immune-precipitated for 5 hr at 4°C using M2-flag agarose beads. Immune-precipitates were analysed by Western blot or by LC-MS/MS (Central Proteomics Facility, Sir William Dunn Pathology School, Oxford University).

#### Functional annotation of the putative TRAPPC2 protein partners

TRAPPC2-FLAG IP samples were compared with the proteins immunoprecipitated in the control IP (refer to Supplementary Table S1): 427 proteins were specifically immunoprecipitated in TRAPPC2-FLAG samples. Moreover, 113 were enriched in the TRAPPC2-FLAG IP samples versus the control IP (ratio >2). Functional Annotation analysis (Dennis et al., 2003; Huang et al., 2009) was used to group the putative TRAPPC2 protein partners according to Molecular Function (MF). The DAVID online tool (DAVID Bioinformatics Resources 6.7) was used restricting the output to all MF terms (MF_FAT). Of note, we found that 251 out of the total 539 proteins were annotated as RNA-binding proteins (GO:0003723) (Ashburner et al., 2000).

#### Digitonin Assay

HeLa cells were exposed to SA or treated with Dinaciclib (10 μM). After treatment, cells were washed twice with permeabilization buffer (25mM HEPES, 125 mM CH3COOK, 2.5 mM Mg(CH3COO)2, 5 mM EGTA, 1mM DTT) and then permeabilized in PB buffer supplemented with 30 μg/mL of digitonin plus the indicated antibody to visualize permeabilized cells. To analyze Sec24C association with SGs, digitonin was left for 6 min at room temperature. Cells were then washed three times with PB buffer and left in PB buffer for 10 min at RT. Finally, samples were fixed with 4% PFA and processed for IF. To analyse Sec24C association at ERES in the presence of Dinaciclib, cells were fixed for 2 min after digitonin permeabilization.

#### High content screening

HeLa cells were plated in 384-well culture plates. The day after, a kinase inhibitor library, purchased from Selleckchem, was dispensed using a liquid-handler system (Hamilton) to give a final concentration of 10 μM. 150 minutes after drug administration, cells were treated with sodium arsenite (300 μM) for 30 min at 37°C in the presence of compounds. Finally, samples were fixed for 10 min at room temperature by adding 1 volume of 4% PFA (paraformaldehyde in PBS) to the growth medium and stained with the appropriate antibodies.

For image acquisition, at least 25 images per field were acquired per well of the 384-well plate using confocal automated microscopy (Opera high content system; Perkin-Elmer). A dedicated script was developed to perform the analysis of Sec24C localization on the different images (Harmony and Acapella software; Perkin-Elmer). The script calculated the co-localization value of Sec24C with the SG marker (G3BP). The results were normalized using positive (mock cells exposed to SA) control samples in the same plate.

P-values were calculated on the basis of mean values from three independent wells. The data are represented as a percentage of Sec24C recruitment in the control cells (100%) using Excel (Microsoft) and Prism software (GraphPad software).

#### Cell cycle analysis by flow cytometry

HeLa cells were harvested and resuspended in PBS. For the fixation, a 9-fold volume of 70% ethanol was added and incubated a 4°C for at least 1hr. Next, cells were centrifuged, washed in PBS and resuspended in PBS containing RNase A 0.1 mg/ml. After incubation for 1hr at 37°C, propidium iodide was added to a final concentration of 10 μg/ml and samples were analyzed in an Accuri C6 flow cytometer.

#### Cell proliferation analysis by High Content Imaging

Cell cycle analysis by high content imaging was performed using the Click-iT Plus EdU Alexa Fluor 488 Imaging Kit (Life Technologies) according to the manufacturer’s instructions. Images were acquired with an automated confocal microscopy (Opera System, Perkin-Elmer) and analyzed through Columbus Image Data Storage and Analysis System (Perkin-Elmer). Nuclear intensity of EdU (5-ethynyl-2′-deoxyuridine, a nucleoside analog of thymidine) in EdU-positive nuclei (S-phase cells) was used as a measure of DNA replication rate.

#### Cell death assay

HeLa cells were pre-treated with CDK inhibitors (SNS-032, Flavopiridol and Dinaciclib) for 150 min and exposed to SA (500 μM, 1 h). Subsequently, the stress stimulus was washed out and cells left to recover for 16 h. To stain dead cells, a fluorophore dye cocktail containing the cell-permeant nuclear dye Hoechst 33342 and the cell-impermeant nuclear dye BOBO ™-3 that selectively labels dying cells (Invitrogen, H3570, B3586), was added to the growth media and incubated for 45 min (37°C, 5% CO2). Cells were analyzed at automated microscopy (Operetta high content system; Perkin-Elmer) using a customized script to calculate the ratio between BOBO ™-3 positive nuclei and total cells.

#### Puromycin Assay

The puromycin (PMY) assay was modified from David et al., JCB 2012 (David et al., 2012). In brief, mock, TRAPPC2-KD or TRAPPC3-KD HeLa cells were exposed to SA (500 μM, 30 min) in DMEM 10% FCS. Cells were washed three times in DMEM 1X and incubated with 9 μM PMY in DMEM for 5 min at 37°C. Samples were lysed in RIPA buffer and processed for Western blot analysis with the anti-puromycin antibody.

#### Transport assay

VSVG-mEOS2-2XUVR8 was a gift from Matthew Kennedy (AddGene plasmid #49803). HeLa cells were transfected with the plasmid for 16 h and treated with SA, CHX and ISRIB for the indicated times. A UV-A lamp was used to illuminate samples (4 pulses, 15 sec each). After the light pulses, cells were left for 10 min at 37°C, then fixed with a volume of 4% PFA and processed for immunofluorescence. The PC-I transport assay was performed in human fibroblasts as previously described (Venditti et al., 2012). For our purposes, cells were treated with SA (300 μM) for 120 min at 40°C and analyzed 10 min after the temperature switch (40 to 32°C). Cells were then fixed and stained with appropriate antibodies.

#### Electron microscopy

EM samples were prepared as previously described (D’Angelo et al., 2007). Briefly, cells were fixed by adding to the culture medium the same volume of a mixture of PHEM buffer (10 mM EGTA, 2 mM MgCl_2_, 60 mM PIPES, 25 mM HEPES, pH 6.9), 4% paraformaldehyde, 2% glutaraldehyde for 2 h, and then stored in storage solution (PHEM buffer, 0.5% paraformaldehyde) overnight. After washing with 0.15 M glycine buffer in PBS, the cells were scraped and pelleted by centrifugation, embedded in 10% gelatin, cooled on ice, and cut into 0.5 mm blocks. The blocks were infused with 2.3 M sucrose and trimmed with an ultramicrotome (Leica Ultracut R) at −120°C using a dry diamond knife.

#### Quantitative real-time PCR

Real-time quantitative PCR (qRT-PCR) was carried out with the LightCycler 480 SYBR Green I mix (Roche) using the Light Cycler 480 II detection system (Roche) with the following conditions: 95 °C, 10 min; (95 °C, 10 s; 60 °C, 10 s; 72 °C, 15 s) × 45 cycles. For expression studies the qRT-PCR results were normalized against an internal control (HPRT1).

## Acknowledgements

We thank Andrea Ballabio, Carmine Settembre, Leandro Raul Soria, Maria Chiara Masone, Rossella Venditti and Graciana Diez Roux for helpful discussion. MADM acknowledges the support of Telethon (grant TGM11CB1), the Italian Association for Cancer Research (AIRC, grant IG2013_14761), European Research Council Advanced Investigator grant no. 670881 (SYSMET).

## Author contributions

F.Z. and M.A.D.M. conceived the work. F.Z. planned and analyzed most of the experiments, C.W. performed experiments on yeast, G.D.T. performed gel filtration and Sec31-IP, M.S. provided technical support, P.P. and M.P. performed MS/MS analysis, D.D. helped with cell culture, S.P.V. performed cell proliferation and flow cytometry assays, R.D.C. and M.F. analyzed MS/MS data, A.R. helped in performing cell death assay, M.A.S. provided podocytes cell line, E.P. performed E.M. analyses, L.G. designed the script for high-content analysis, F.Z. and M.A.D.M. conceptualized the work and strategy and wrote the manuscript.

---

**Figure 1—figure supplement 1.**
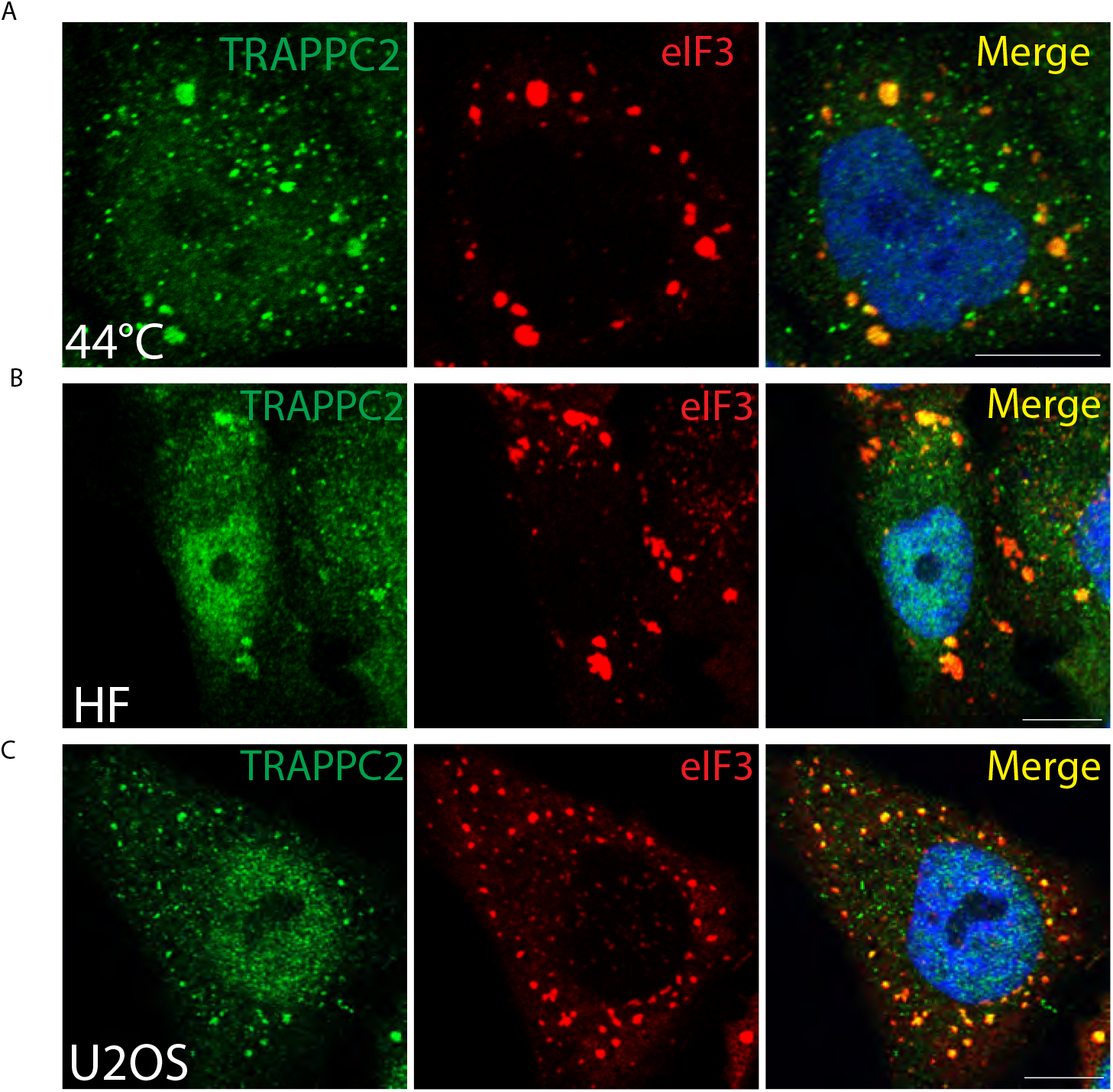
Re-localization of TRAPP to SGs is independent of stress stimulus and cell type. **(A)** HeLa cells exposed to heat shock (44°C, 45 min), **(B)** human fibroblasts treated with SA (500 μM, 180 min), **(C)** U2OS cells treated with SA (300 μM, 60 min), stained for eIF3 and TRAPPC2. Scale bar, 10 μm.

**Figure 1—figure supplement 2.**
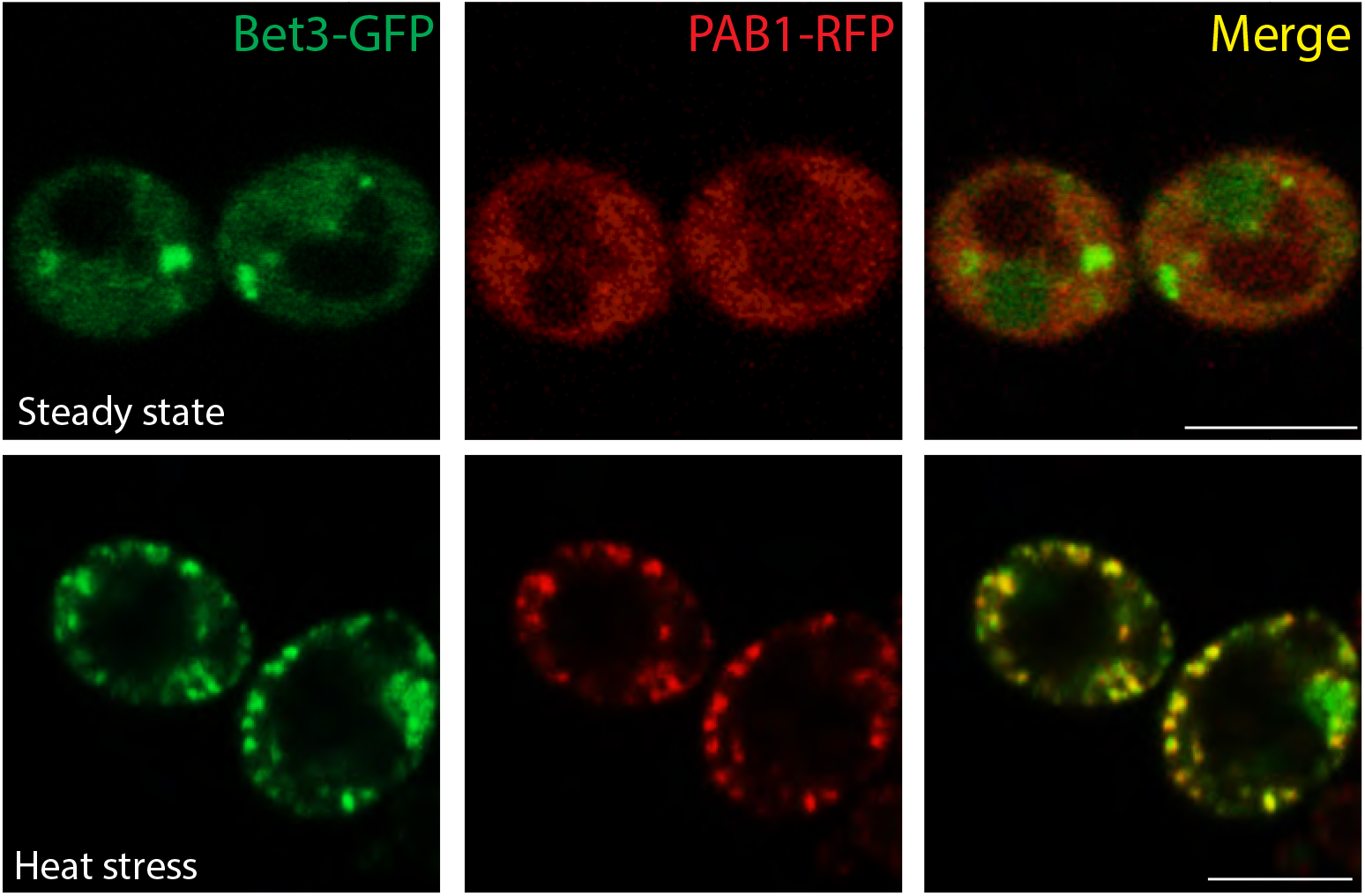
Stress-induced localization of Bet3 (TRAPPC3) to SGs in yeast cells. Cells co-expressing Bet3-GFP and the stress granule marker RFP-PAB1 were exposed to heat shock (46°C, 10 min) and imaged. Scale bar, 5μm. **Figure 1—supplement video 1. Figure 1—supplement video 2.**

**Figure 1—figure supplement 3.**
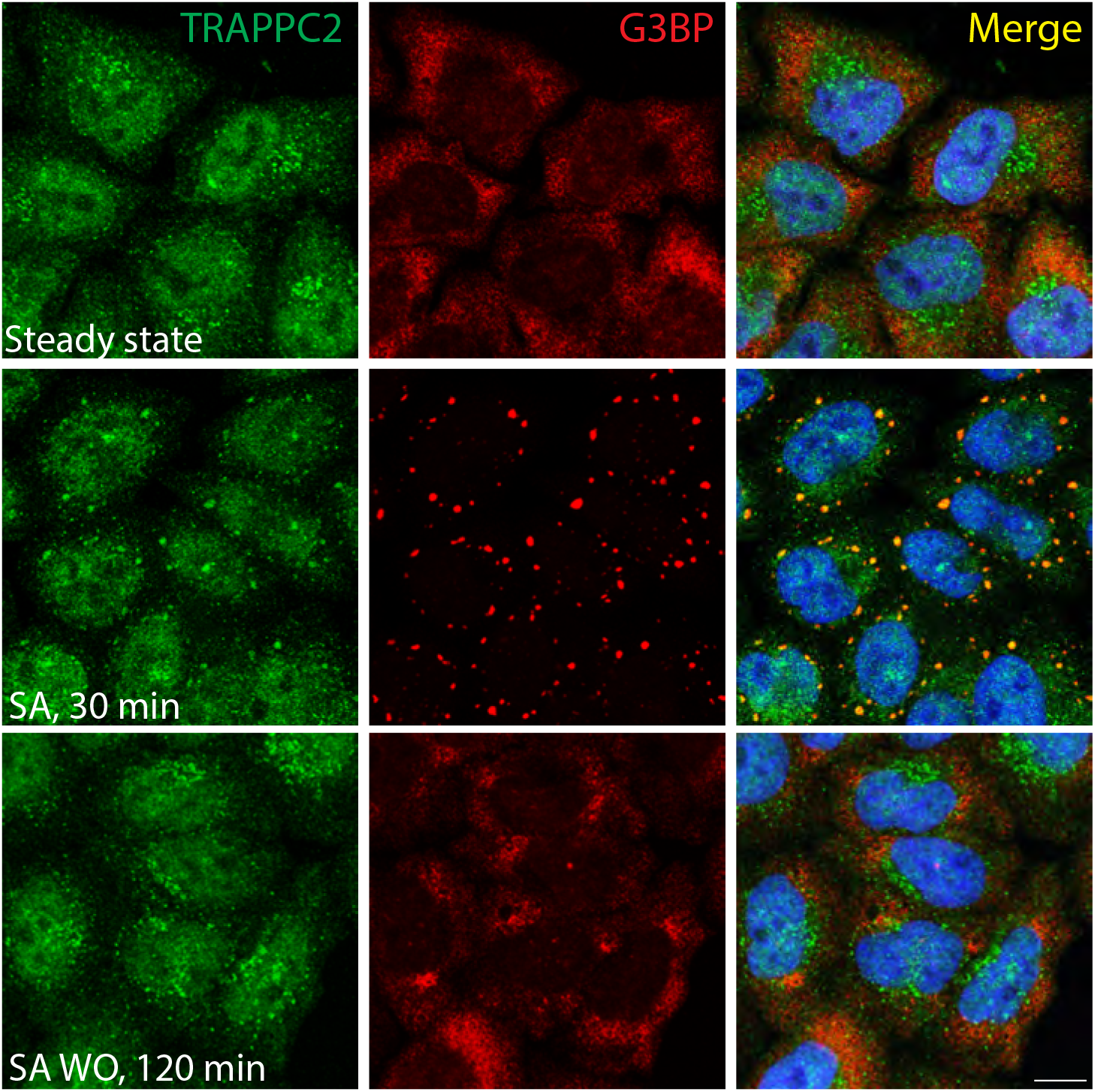
Association of TRAPP with SGs is reversible after SG disassembly. Cells were treated with SA for 30 min, the SA was washed out, and cells were left to recover for 120 min and processed for staining. Scale bar, 10 μm.

**Figure 2—figure supplement 1.**
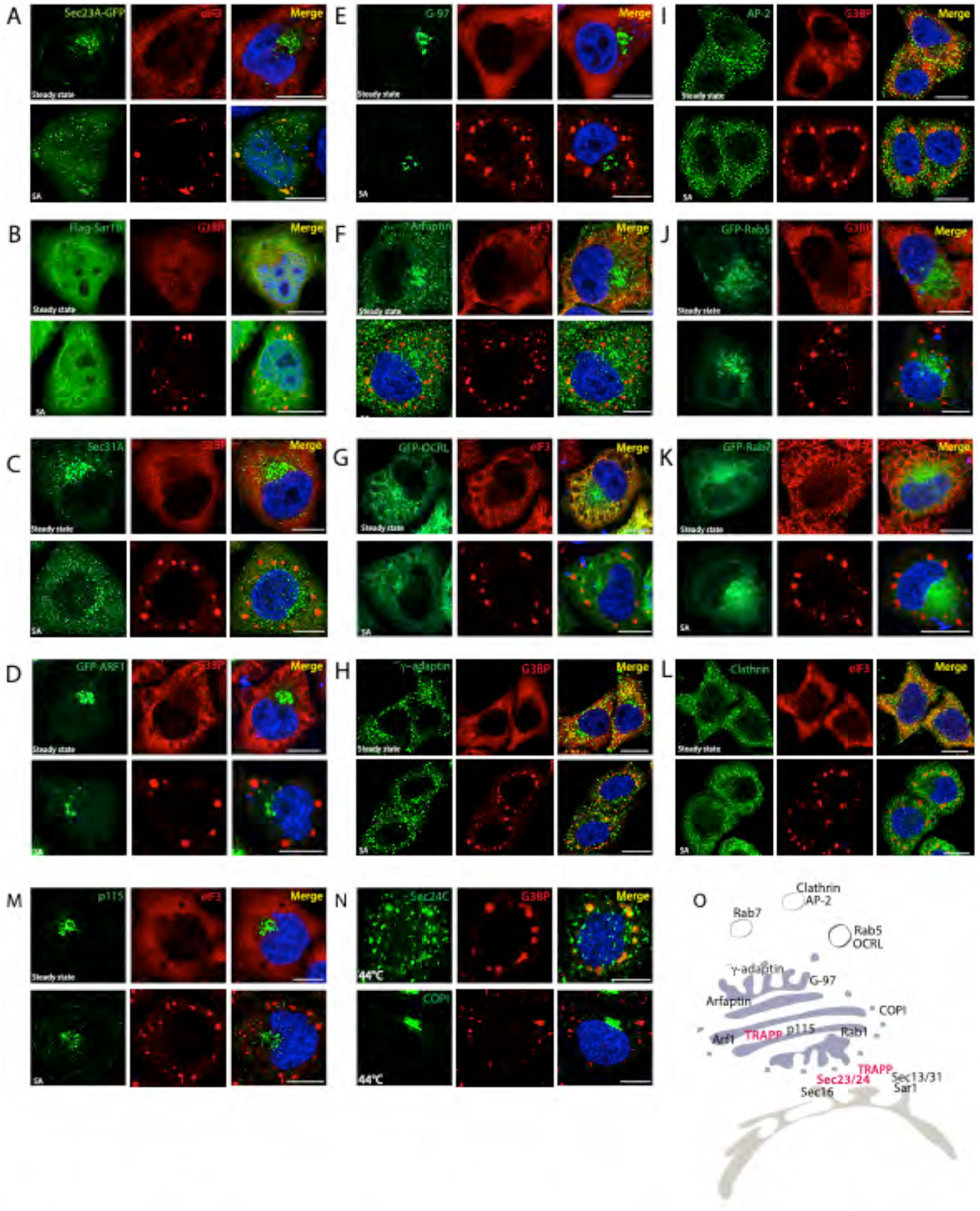
Sec24C and Sec23, but not components of other coat complexes or other cytosolic proteins associated with the exocytic and endocytic pathways, translocate to SGs. **(A-M)** HeLa cells at steady state or treated with SA for 30 min were visualized by fluorescence microscopy using antibodies against the endogenous proteins or Flag-tagged proteins, or visualizing transfected GFP-tagged proteins, and co-stained for eIF3 or G3BP, as indicated. **(N)** HeLa cells exposed to heat shock (44°C, 45 min) were stained for Sec24C or COPI, and G3BP. Blue, DAPI. Scale bar, 10 μm. **(O)** Schematic diagram showing the cellular location of the tested proteins that associate (red) or not (black) with SGs.

**Figure 2—figure supplement 2.**
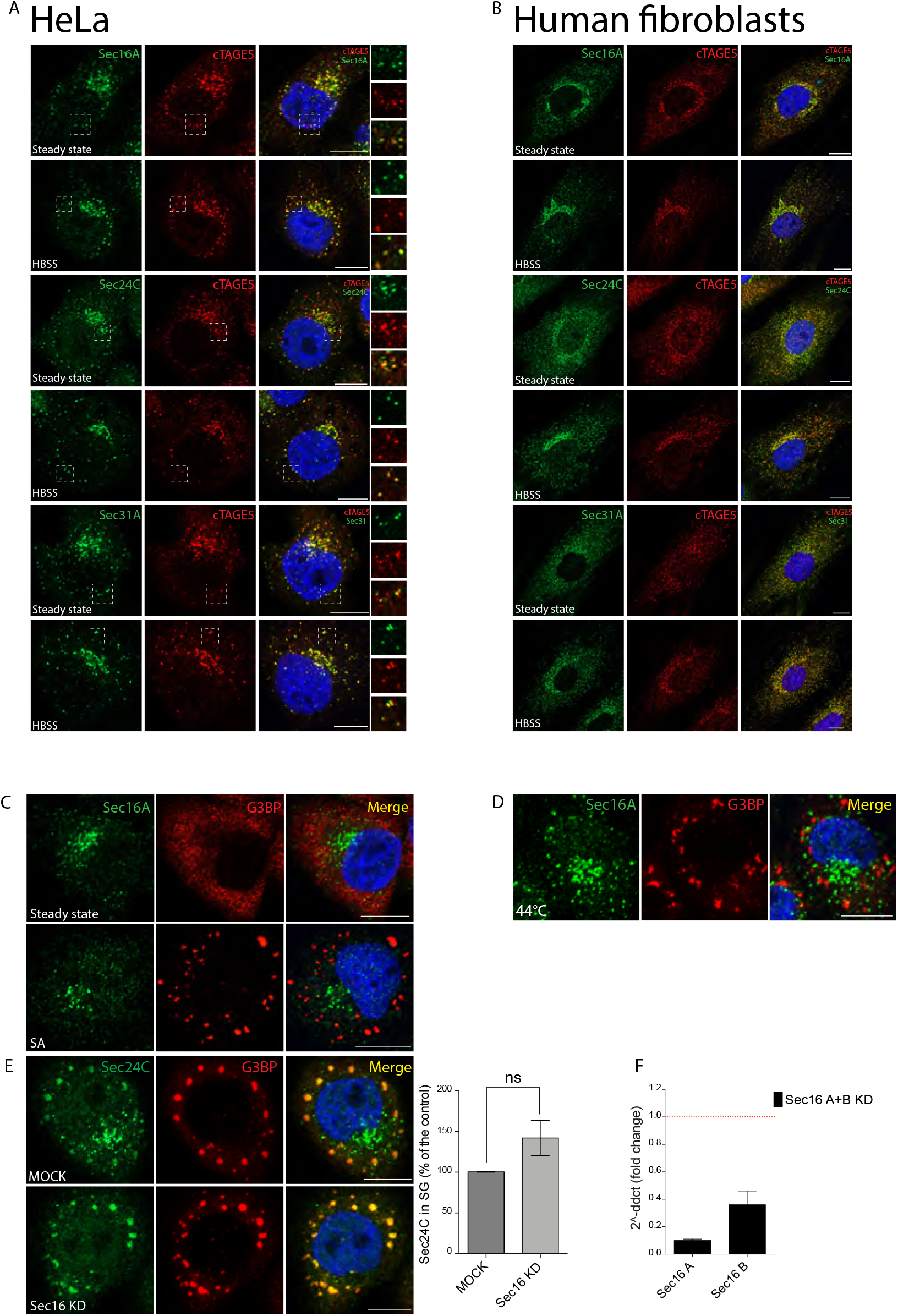
Sec bodies do not form in mammalian cells exposed to aa starvation and Sec16 is not required for SG assembly in response to oxidative stress or for the recruitment of COPII to SGs. **(A)** HeLa cells and human fibroblasts **(B)** were aa starved in HBSS for 8 hr and stained for cTAGE5 (red) to visualize ERES, and for Sec16 or the indicated COPII components (Sec24C and Sec31A)(green). **(C)** Cells were treated with SA (300 μM for 30 min) and co-stained for Sec16 and G3BP as an SG marker. **(D)** Cells exposed to heat shock were processed as described in (A). **(E)** Representative images of Sec24C localization in Sec16A+B KD (Sec16 KD) cells treated with SA. Cells were fixed and visualized by fluorescence microscopy using anti-Sec24C Ab, anti-G3BP Ab, and DAPI (blue). Quantification of TRAPPC2 redistribution to SGs calculated as the ratio between Sec24C (mean intensity) in SG puncta and cytosolic Sec24C. Mean ± s.e.m., n = 40-60 cells per experiment, N = 3. ns: not significant. Scale bar 10 μm. **(F)** Sec16A and B mRNA levels in Sec16A+B KD cells were evaluated by q-PCR and expressed as fold change with respect to the MOCK. Dashed red line: MOCK.

**Figure 2—figure supplement 3.**
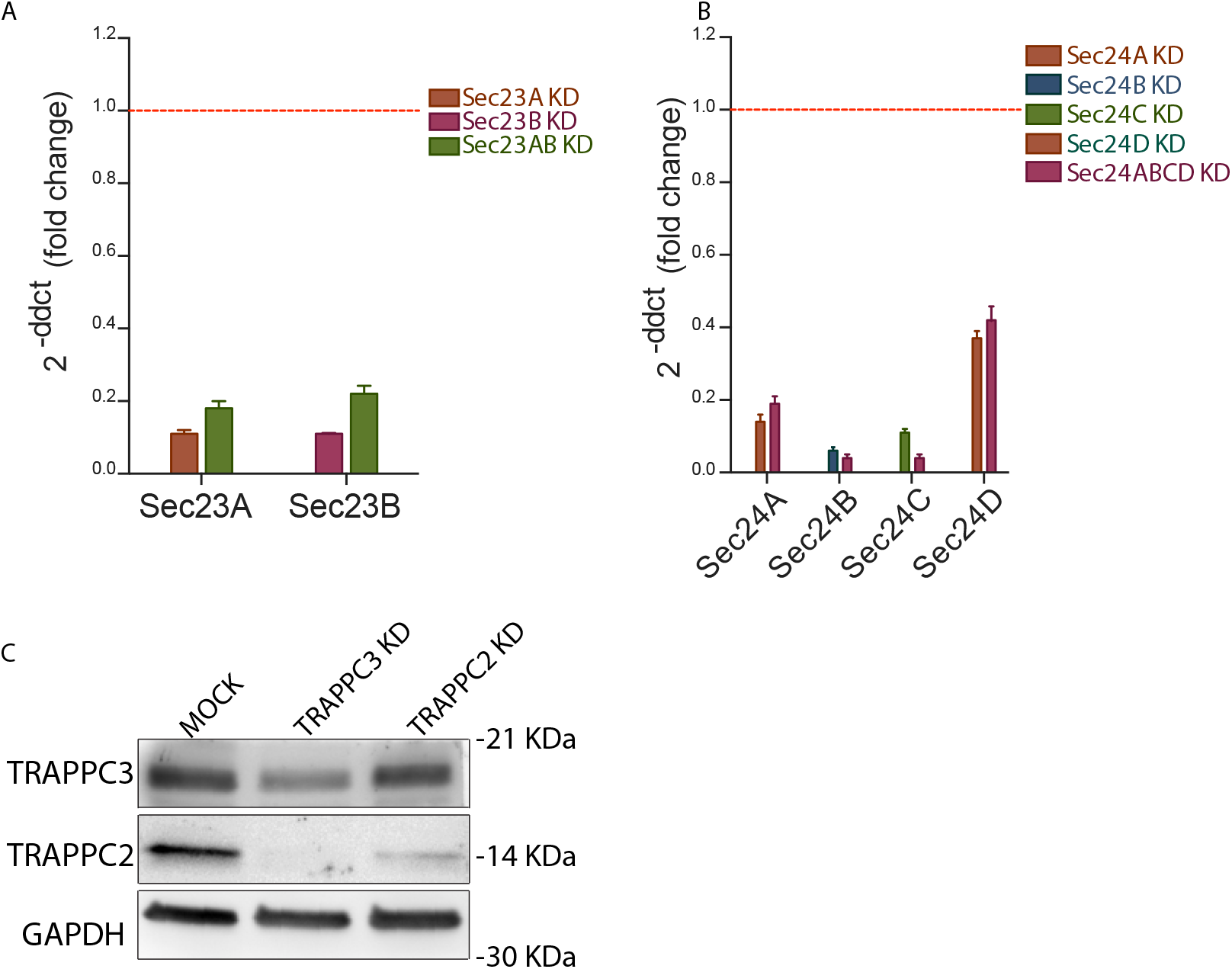
Evaluation of Sec23, Sec24, TRAPPC2 and TRAPPC3 knock down efficiencies. **(A)** Sec23A and Sec23B mRNA levels were evaluated by q-PCR and expressed as fold change with respect to the MOCK. Dashed red line: MOCK. **(B)** Sec24A, B, C and D mRNA levels were evaluated as described in (A). **(C)** Representative Western blot of HeLa cell extracts depleted for TRAPPC2 and TRAPPC3; GAPDH was used as loading control.

**Figure 3—figure supplement 1.**
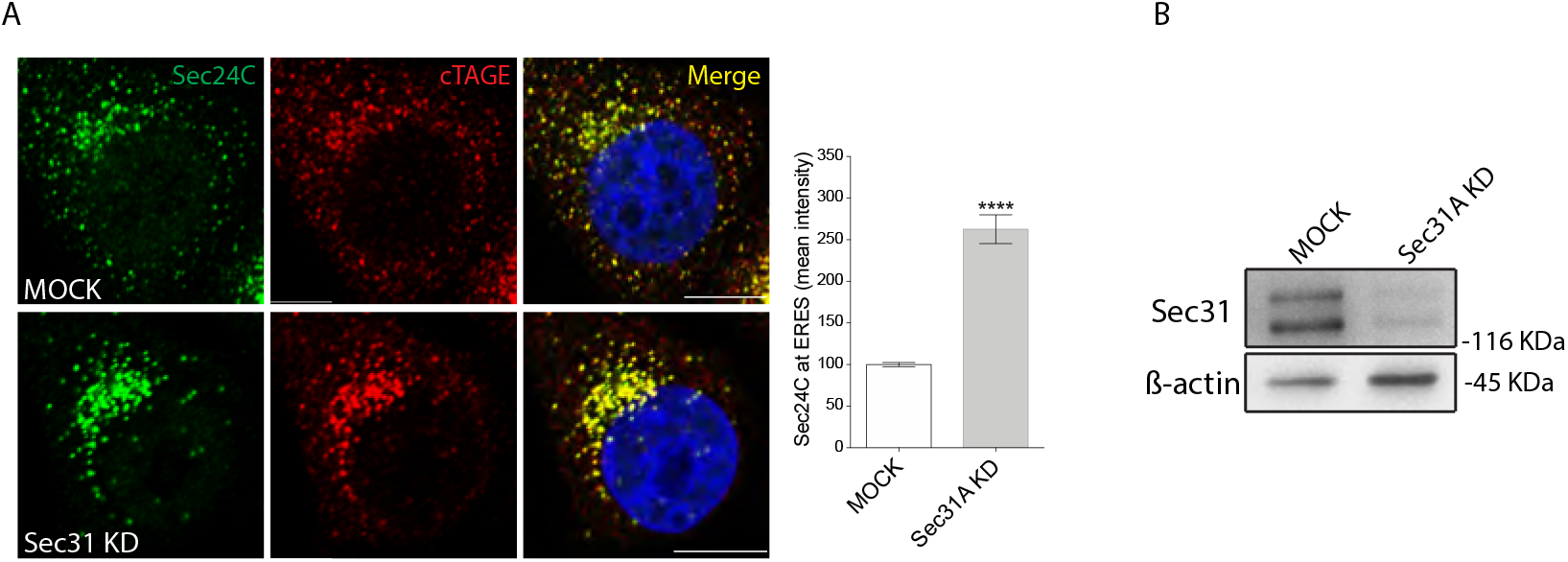
Effect of Sec31 depletion on Sec24 localization at ERES. **(A)** Cells were mock-treated or KD for Sec31 and then immunostained for Sec24C and cTAGE5 to stain ERES. Scale bar 10μm. The graph shows the mean intensity of Sec24C at ERES normalized for cytosolic signal as a percentage of control conditions. Scale bar 10 μm. **(B)** Western blot analysis of Sec31A KD cells. β-actin was used as loading control.

**Figure 4—figure supplement 1.**
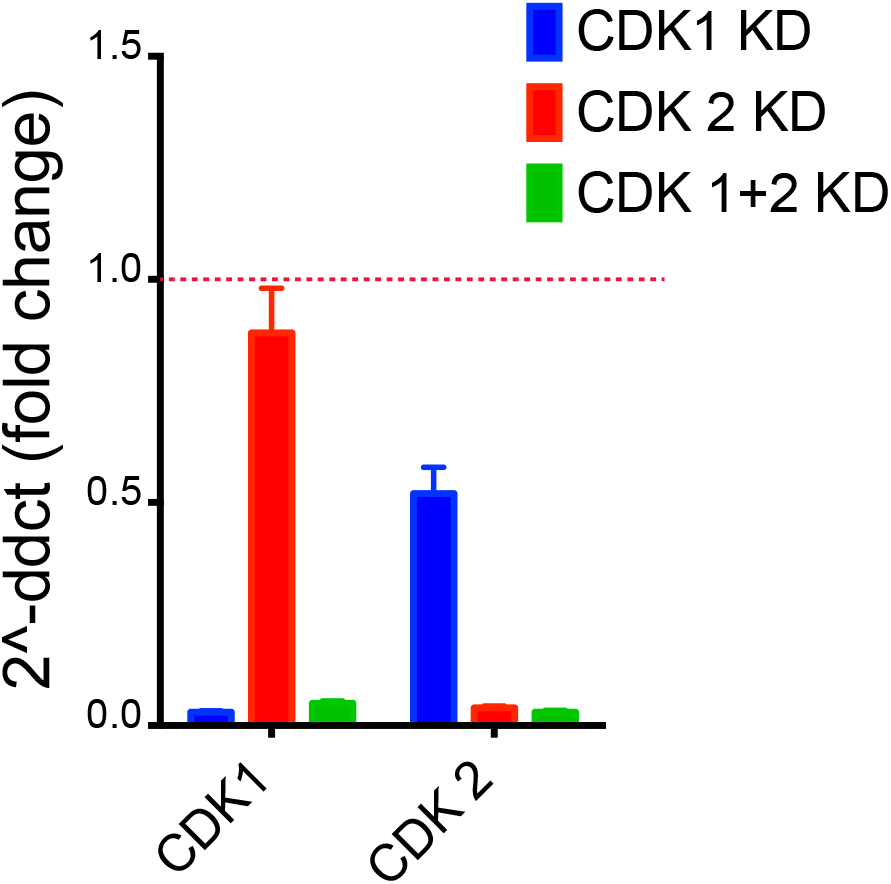
Evaluation of CDK1 and CDK2 knock down efficiency. CDK1 and CDK2 mRNA levels were evaluated by q-PCR and expressed as fold change with respect to the MOCK. Dashed red line: MOCK.

**Figure 6—figure supplement 1.**
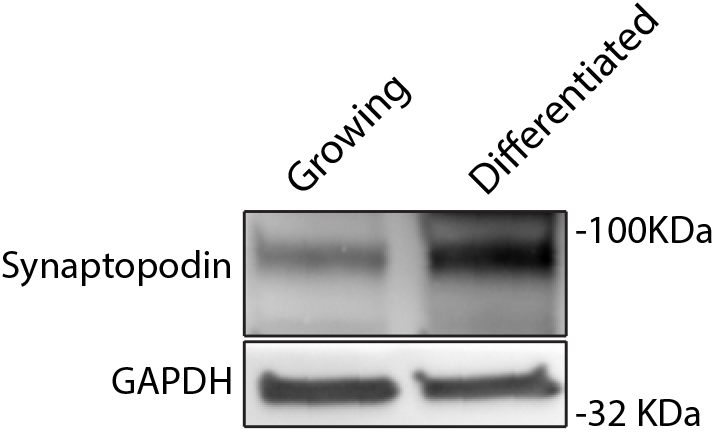
Differentiation of podocytes. Western blot of growing and differentiated podocytes. Synaptopodin was used as a differentiation marker, GAPDH as a loading control.

**Figure 7—figure supplement 1.**
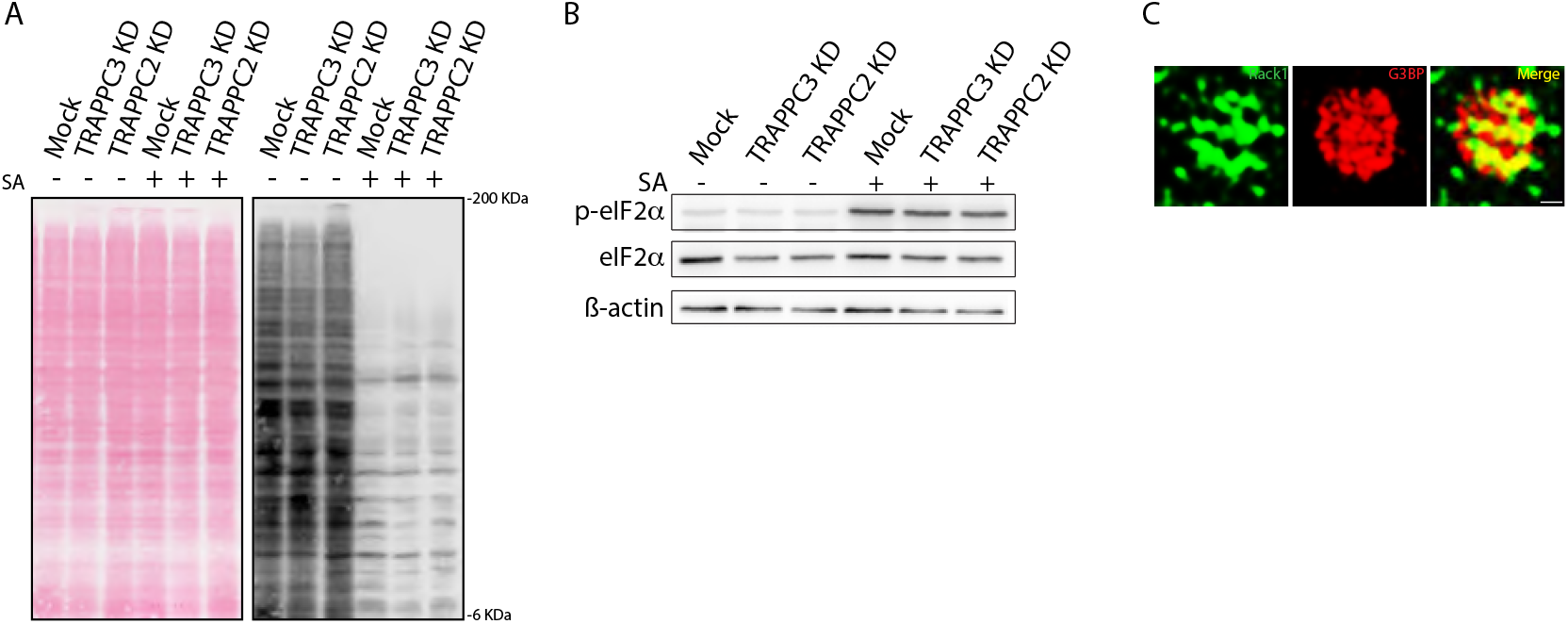
TRAPP depletion does not affect protein translation inhibition caused by SA treatment. **(A)** Puromycin assay (see Materials and methods) of MOCK, TRAPPC3 and TRAPPC2-KD HeLa cells at steady state and upon SA treatment. **(B)** Evaluation of phosphorylated eIF2a in TRAPPC3 and TRAPPC2 depleted cells exposed to SA. ß-actin was used as a loading control. **(C)** Structured Illumination Microscopy (SIM)-Super resolution (SR) images of endogenous RACK1 and G3BP. Scale bar 500 nm.

**Figure 8—figure supplement 1.**
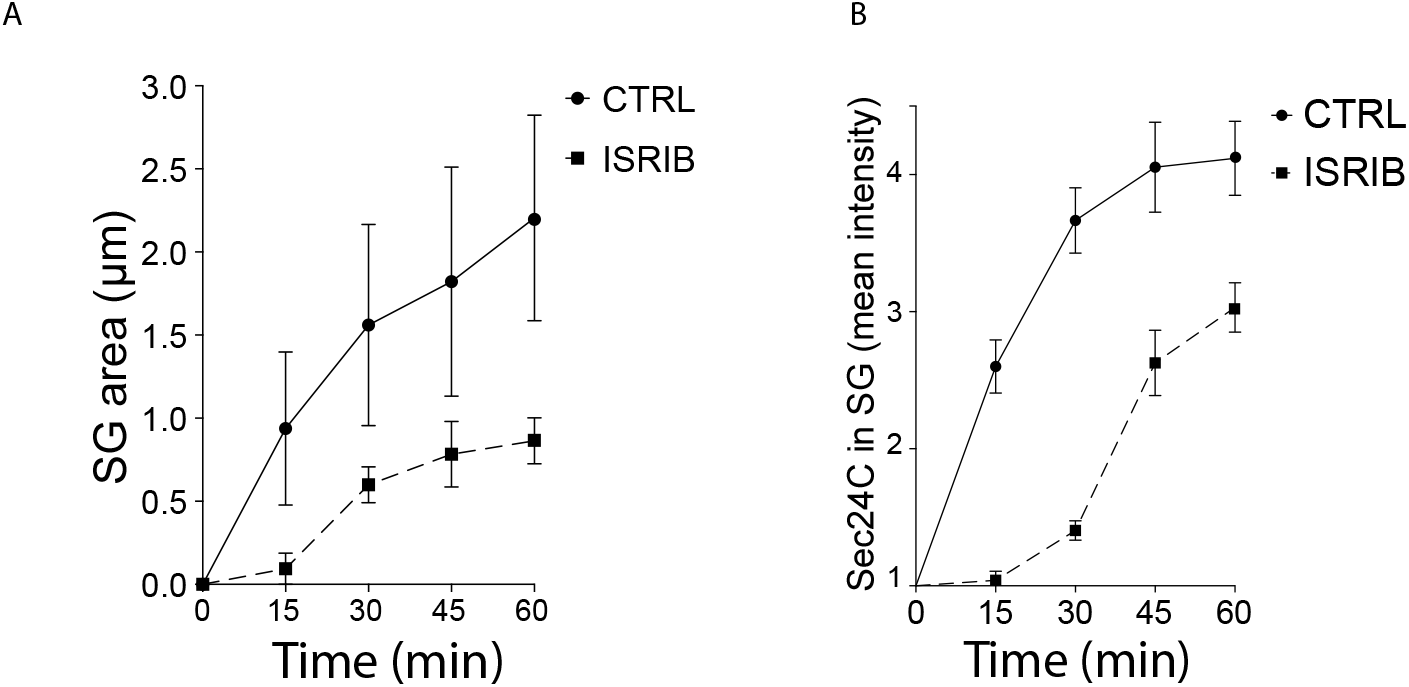
ISRIB delays SG nucleation and prevents Sec24C relocalization to SGs. **(A)** HeLa cells were exposed to SA (200 μM) for the indicated time, alone or in combination with ISRIB and SGs were analyzed. Quantification of SG area (μm) of three independent experiment Mean ± s.e.m. **(B)** Graph, Quantification of Sec24C localization at SGs over time (the ratio between Sec24C (mean fluorescence intensity) in SG puncta and cytosolic Sec24C). Mean ± SD. n = 60-80 cells per experiment, N = 3.

**Figure 9—figure supplement 1.**
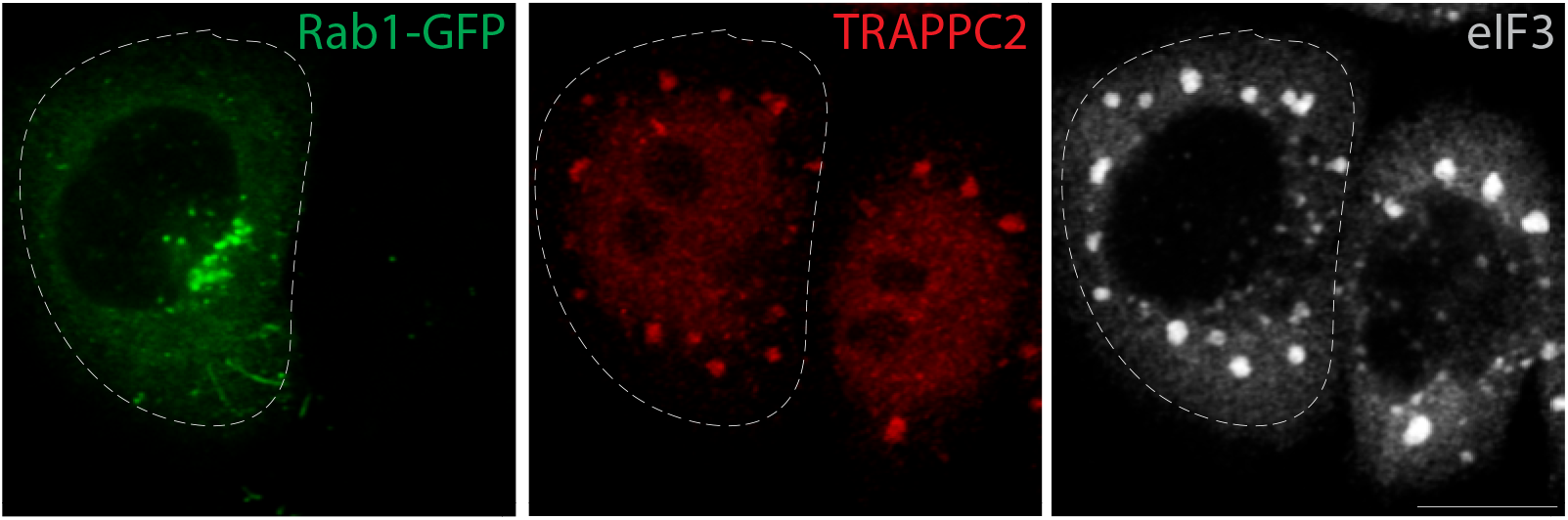
TRAPPC2 migrates to SGs in Rab1 overexpressing cells. HeLa cells overexpressing GFP-Rab1B-WT were treated with SA (300 μM 30 min) and stained for TRAPPC2 (in red) and eIF3 (in gray). Dashed white line: Overexpressing cells.

**Table S1.**
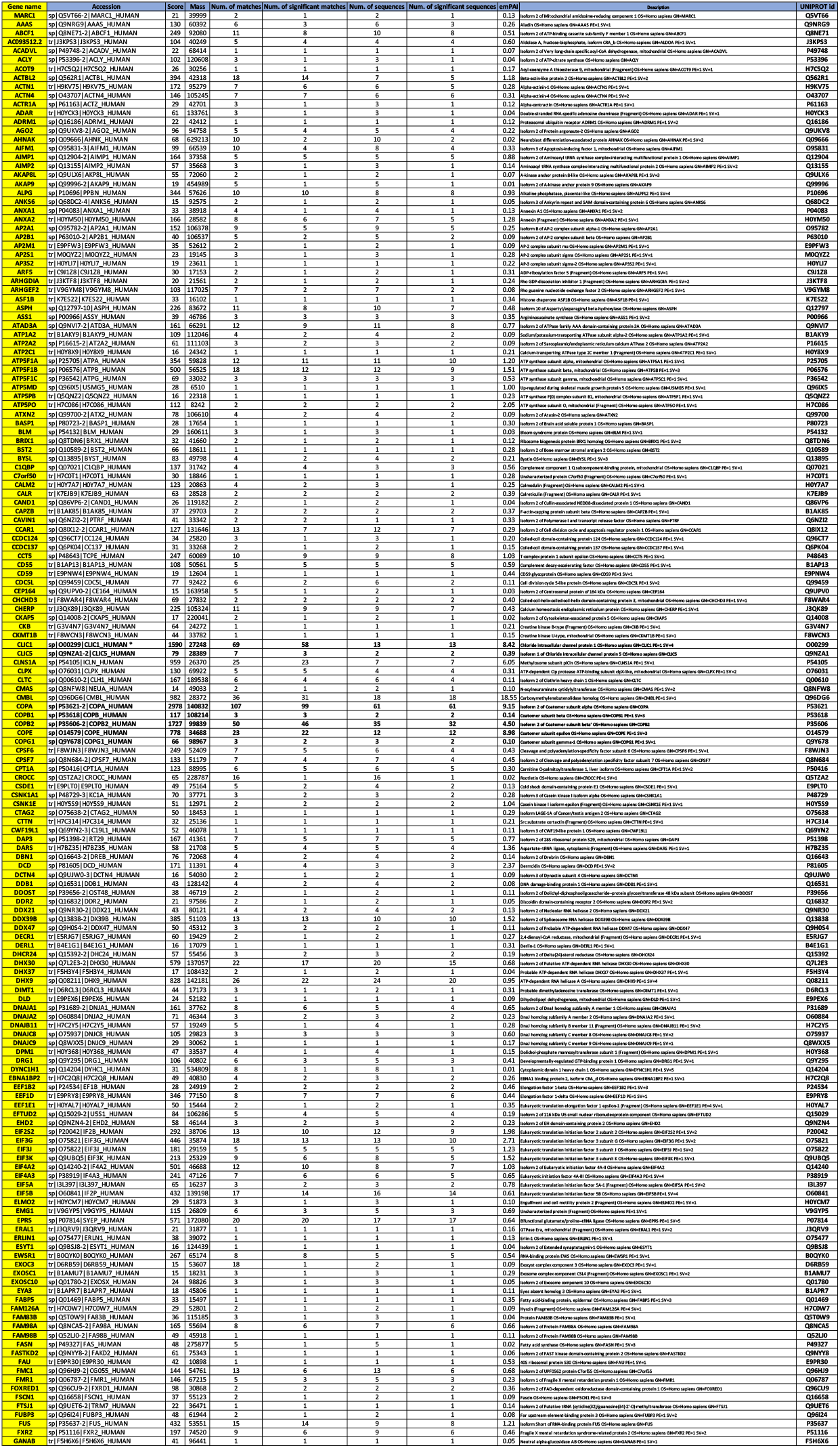
TRAPPC2 interactors.

**Table S2.**
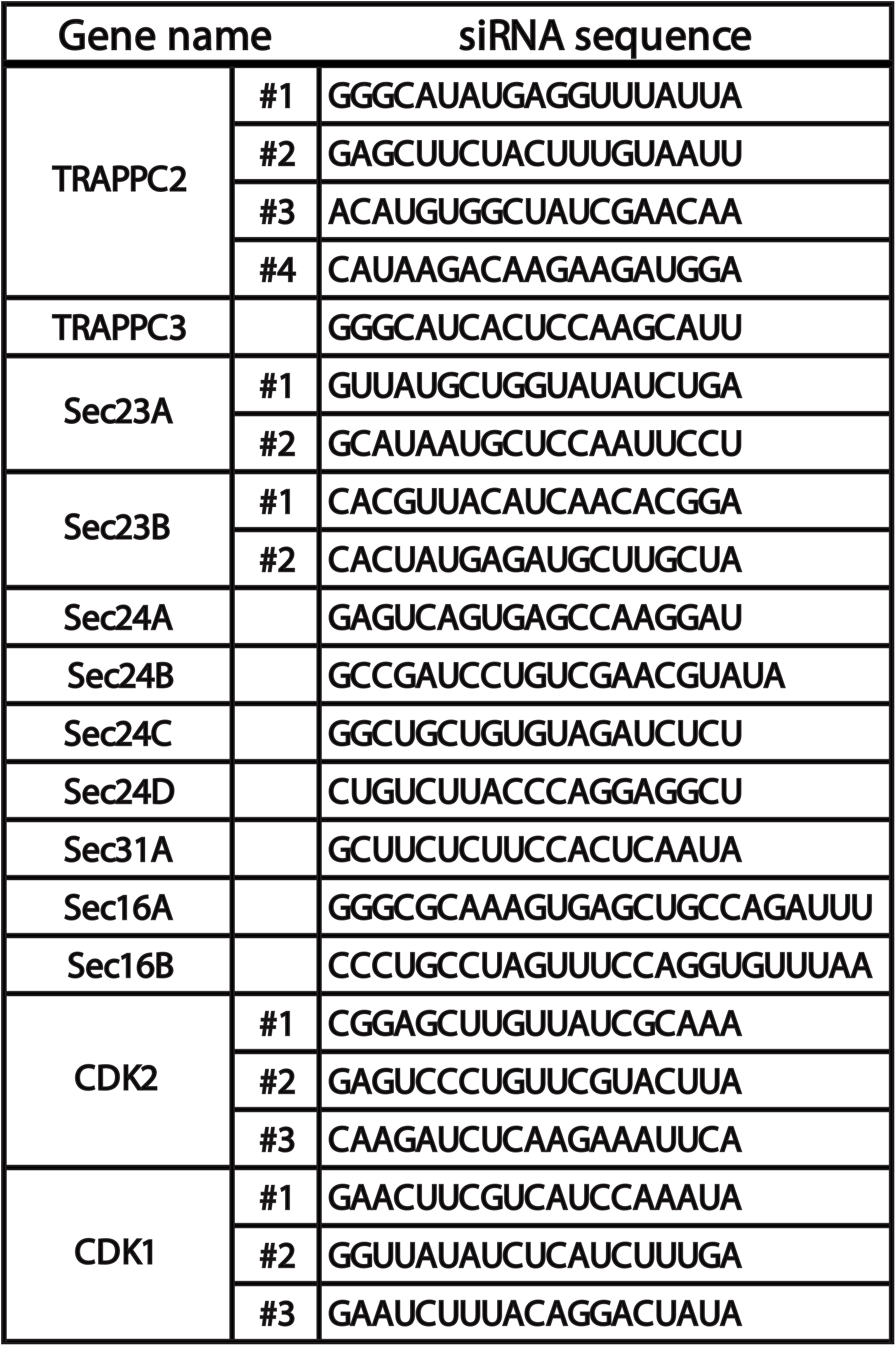
Small interfering RNA oligonucleotides used in the study.

## References

Aguilera-Gomez, A., M. Zacharogianni, M.M. van Oorschot, H. Genau, R. Grond, T. Veenendaal, K.S. Sinsimer, E.R. Gavis, C. Behrends, and C. Rabouille. 2017. Phospho-Rasputin Stabilization by Sec16 Is Required for Stress Granule Formation upon Amino Acid Starvation. Cell Reports. 20:935–948.

Anderson P, Kedersha N. 2008. Stress granules: the Tao of RNA triage. Trends Biochem Sci 33:141–150. doi:10.1016/j.tibs.2007.12.003

Anderson P, Kedersha N. 2002. Stressful initiations. J Cell Sci 115:3227–3234.

Arimoto K, Fukuda H, Imajoh-Ohmi S, Saito H, Takekawa M. 2008. Formation of stress granules inhibits apoptosis by suppressing stress-responsive MAPK pathways. Nat Cell Biol 10:1324–1332. doi:10.1038/ncb1791

Asghar U, Witkiewicz AK, Turner NC, Knudsen ES. 2015. The history and future of targeting cyclin-dependent kinases in cancer therapy. Nat Rev Drug Discov 14:130–146. doi:10.1038/nrd4504

Ashburner M, Ball CA, Blake JA, Botstein D, Butler H, Cherry JM, Davis AP, Dolinski K, Dwight SS, Eppig JT, Harris MA, Hill DP, Issel-Tarver L, Kasarskis A, Lewis S, Matese JC, Richardson JE, Ringwald M, Rubin GM, Sherlock G. 2000. Gene ontology: tool for the unification of biology. The Gene Ontology Consortium. Nat Genet 25:25–29. doi:10.1038/75556

Aulas A, Fay MM, Szaflarski W, Kedersha N, Anderson P, Ivanov P. 2017. Methods to Classify Cytoplasmic Foci as Mammalian Stress Granules. J Vis Exp JoVE. doi:10.3791/55656

Ballock RT, O’keefe RJ. 2003. The Biology Of The Growth Plate. J Bone Jt Surg-Am Vol 85:715–726.

Bi X, Mancias JD, Goldberg J. 2007. Insights into COPII Coat Nucleation from the Structure of Sec23·Sar1 Complexed with the Active Fragment of Sec31. Dev Cell 13:635–645. doi:10.1016/j.devcel.2007.10.006

Catara G, Grimaldi G, Schembri L, Spano D, Turacchio G, Lo Monte M, Beccari AR, Valente C, Corda D. 2017. PARP1-produced poly-ADP-ribose causes the PARP12 translocation to stress granules and impairment of Golgi complex functions. Sci Rep 7:14035. doi:10.1038/s41598-017-14156-8

Chen D, Gibson ES, Kennedy MJ. 2013. A light-triggered protein secretion system. J Cell Biol 201:631–640. doi:10.1083/jcb.201210119

David A, Dolan BP, Hickman HD, Knowlton JJ, Clavarino G, Pierre P, Bennink JR, Yewdell JW. 2012. Nuclear translation visualized by ribosome-bound nascent chain puromycylation. J Cell Biol 197:45–57. doi:10.1083/jcb.201112145

D’Arcangelo JG, Stahmer KR, Miller EA. 2013. Vesicle-mediated export from the ER: COPII coat function and regulation. Biochim Biophys Acta 1833:2464–2472. doi:10.1016/j.bbamcr.2013.02.003

De Leo MG, Staiano L, Vicinanza M, Luciani A, Carissimo A, Mutarelli M, Di Campli A, Polishchuk E, Di Tullio G, Morra V, Levtchenko E, Oltrabella F, Starborg T, Santoro M, Di Bernardo D, Devuyst O, Lowe M, Medina DL, Ballabio A, De Matteis MA. 2016. Autophagosome-lysosome fusion triggers a lysosomal response mediated by TLR9 and controlled by OCRL. Nat Cell Biol 18:839–850. doi:10.1038/ncb3386

Dennis G, Sherman BT, Hosack DA, Yang J, Gao W, Lane HC, Lempicki RA. 2003. DAVID: Database for Annotation, Visualization, and Integrated Discovery. Genome Biol 4:P3.

Dephoure N, Zhou C, Villén J, Beausoleil SA, Bakalarski CE, Elledge SJ, Gygi SP. 2008 A quantitative atlas of mitotic phosphorylation. Proc Natl Acad Sci U S A. 105(31): 10762–7

Dubois T, Paléotti O, Mironov AA, Fraisier V, Stradal TEB, De Matteis MA, Franco M, Chavrier P. 2005. Golgi-localized GAP for Cdc42 functions downstream of ARF1 to control Arp2/3 complex and F-actin dynamics. Nat Cell Biol 7:353–364. doi:10.1038/ncb1244

Farhan H, Wendeler MW, Mitrovic S, Fava E, Silberberg Y, Sharan R, Zerial M, Hauri H-P. 2010. MAPK signaling to the early secretory pathway revealed by kinase/phosphatase functional screening. J Cell Biol 189:997–1011. doi:10.1083/jcb.200912082

Franzmann TM, Alberti S. 2019. Protein Phase Separation as a Stress Survival Strategy. Cold Spring Harb Perspect Biol. doi:10.1101/cshperspect.a034058

Ge L, Zhang M, Kenny SJ, Liu D, Maeda M, Saito K, Mathur A, Xu K, Schekman R. 2017. Remodeling of ER-exit sites initiates a membrane supply pathway for autophagosome biogenesis. EMBO Rep 18:1586–1603. doi:10.15252/embr.201744559

Gedeon AK, Colley A, Jamieson R, Thompson EM, Rogers J, Sillence D, Tiller GE, Mulley JC, Gécz J. 1999. Identification of the gene (SEDL) causing X-linked spondyloepiphyseal dysplasia tarda. Nat Genet 22:400–404. doi:10.1038/11976

Gorur A, Yuan L, Kenny SJ, Baba S, Xu K, Schekman R. 2017. COPII-coated membranes function as transport carriers of intracellular procollagen I. J Cell Biol 216:1745–1759. doi:10.1083/jcb.201702135

Harripaul R, Vasli N, Mikhailov A, Rafiq MA, Mittal K, Windpassinger C, Sheikh TI, Noor A, Mahmood H, Downey S, Johnson M, Vleuten K, Bell L, Ilyas M, Khan FS, Khan V, Moradi M, Ayaz M, Naeem F, Heidari A, Ahmed I, Ghadami S, Agha Z, Zeinali S, Qamar R, Mozhdehipanah H, John P, Mir A, Ansar M, French L, Ayub M, Vincent JB. 2018. Mapping autosomal recessive intellectual disability: combined microarray and exome sequencing identifies 26 novel candidate genes in 192 consanguineous families. Mol Psychiatry 23:973–984. doi:10.1038/mp.2017.60

Henrotin YE, Bruckner P, Pujol J-PL. 2003. The role of reactive oxygen species in homeostasis and degradation of cartilage. Osteoarthritis Cartilage 11:747–755.

Holt LJ, Tuch BB, Villén J, Johnson AD, Gygi SP, Morgan DO. 2009. Global analysis of Cdk1 substrate phosphorylation sites provides insights into evolution. Science 325:1682–1686. doi:10.1126/science.1172867

Hu H, Gourguechon S, Wang CC, Li Z. 2016. The G1 Cyclin-dependent Kinase CRK1 in Trypanosoma brucei Regulates Anterograde Protein Transport by Phosphorylating the COPII Subunit Sec31. J Biol Chem 291:15527–15539. doi:10.1074/jbc.M116.715185

Yu S, Satoh A, Pypaert M, Mullen K, Hay JC, Ferro-Novick S. 2006. mBet3p is required for homotypic COPII vesicle tethering in mammalian cells. J Cell Biol 174:359–368. doi:10.1083/jcb.200603044

Huang DW, Sherman BT, Zheng X, Yang J, Imamichi T, Stephens R, Lempicki RA. 2009. Extracting biological meaning from large gene lists with DAVID. Curr Protoc Bioinforma Chapter 13:Unit 13.11. doi:10.1002/0471250953.bi1311s27

Imai K, Hao F, Fujita N, Tsuji Y, Oe Y, Araki Y, Hamasaki M, Noda T, Yoshimori T. 2016. Atg9A trafficking through the recycling endosomes is required for autophagosome formation. J Cell Sci 129:3781–3791. doi:10.1242/jcs.196196

Jain S, Wheeler JR, Walters RW, Agrawal A, Barsic A, Parker R. 2016. ATPase modulated stress granules contain a diverse proteome and substructure. Cell 164:487–498. doi:10.1016/j.cell.2015.12.038

Kapetanovich L, Baughman C, Lee TH. 2005. Nm23H2 Facilitates Coat Protein Complex II Assembly and Endoplasmic Reticulum Export in Mammalian Cells. Mol Biol Cell 16:835–848. doi:10.1091/mbc.E04-09-0785

Khattak NA, Mir A. 2014. Computational analysis of TRAPPC9: candidate gene for autosomal recessive non-syndromic mental retardation. CNS Neurol Disord Drug Targets 13:699–711.

Kim JJ, Lipatova Z, Segev N. 2016. TRAPP Complexes in Secretion and Autophagy. Front Cell Dev Biol 4:20. doi:10.3389/fcell.2016.00020

Koehler K, Milev MP, Prematilake K, Reschke F, Kutzner S, Jühlen R, Landgraf D, Utine E, Hazan F, Diniz G, Schuelke M, Huebner A, Sacher M. 2017. A novel TRAPPC11 mutation in two Turkish families associated with cerebral atrophy, global retardation, scoliosis, achalasia and alacrima. J Med Genet 54:176–185. doi:10.1136/jmedgenet-2016-104108

Koreishi M, Yu S, Oda M, Honjo Y, Satoh A. 2013. CK2 Phosphorylates Sec31 and Regulates ER-To-Golgi Trafficking. PloS One 8:e54382. doi:10.1371/journal.pone.0054382

Lamb CA, Nühlen S, Judith D, Frith D, Snijders AP, Behrends C, Tooze SA. 2016. TBC1D14 regulates autophagy via the TRAPP complex and ATG9 traffic. EMBO J 35:281–301. doi:10.15252/embj.201592695

Li C, Luo X, Zhao S, Siu GK, Liang Y, Chan HC, Satoh A, Yu SS. 2017. COPI-TRAPPII activates Rab18 and regulates its lipid droplet association. EMBO J 36:441–457. doi:10.15252/embj.201694866

Lord C, Bhandari D, Menon S, Ghassemian M, Nycz D, Hay J, Ghosh P, Ferro-Novick S. 2011. Sequential interactions with Sec23 control the direction of vesicle traffic. Nature 473:181–186. doi:10.1038/nature09969

Malumbres M. 2014. Cyclin-dependent kinases. Genome Biol 15:122.

Milev MP, Graziano C, Karall D, Kuper WFE, Al-Deri N, Cordelli DM, Haack TB, Danhauser K, Iuso A, Palombo F, Pippucci T, Prokisch H, Saint-Dic D, Seri M, Stanga D, Cenacchi G, van Gassen KLI, Zschocke J, Fauth C, Mayr JA, Sacher M, van Hasselt PM. 2018. Bi-allelic mutations in TRAPPC2L result in a neurodevelopmental disorder and have an impact on RAB11 in fibroblasts. J Med Genet 55:753–764. doi:10.1136/jmedgenet-2018-105441

Milev MP, Grout ME, Saint-Dic D, Cheng Y-HH, Glass IA, Hale CJ, Hanna DS, Dorschner MO, Prematilake K, Shaag A, Elpeleg O, Sacher M, Doherty D, Edvardson S. 2017. Mutations in TRAPPC12 Manifest in Progressive Childhood Encephalopathy and Golgi Dysfunction. Am J Hum Genet 101:291–299. doi:10.1016/j.ajhg.2017.07.006

Mironov AA, Beznoussenko GV, Nicoziani P, Martella O, Trucco A, Kweon H-S, Giandomenico DD, Polishchuk RS, Fusella A, Lupetti P, Berger EG, Geerts WJC, Koster AJ, Burger KNJ, Luini A. 2001. Small cargo proteins and large aggregates can traverse the Golgi by a common mechanism without leaving the lumen of cisternae. J Cell Biol 155:1225–1238. doi:10.1083/jcb.200108073

Mohamoud HS, Ahmed S, Jelani M, Alrayes N, Childs K, Vadgama N, Almramhi MM, Al-Aama JY, Goodbourn S, Nasir J. 2018. A missense mutation in TRAPPC6A leads to build-up of the protein, in patients with a neurodevelopmental syndrome and dysmorphic features. Sci Rep 8:2053. doi:10.1038/s41598-018-20658-w

Moujalled D, James JL, Yang S, Zhang K, Duncan C, Moujalled DM, Parker SJ, Caragounis A, Lidgerwood G, Turner BJ, Atkin JD, Grubman A, Liddell JR, Olsen JV^1^, Blagoev B, Gnad F, Macek B, Kumar C, Mortensen P, Mann M 2006 Global, in vivo, and site-specific phosphorylation dynamics in signaling networks Cell. 2006; 127(3):635–48

Proepper C, Boeckers TM, Kanninen KM, Blair I, Crouch PJ, White AR. 2015. Phosphorylation of hnRNP K by cyclin-dependent kinase 2 controls cytosolic accumulation of TDP-43. Hum Mol Genet 24:1655–1669. doi:10.1093/hmg/ddu578

Palmer KJ, Konkel JE, Stephens DJ. 2005. PCTAIRE protein kinases interact directly with the COPII complex and modulate secretory cargo transport. J Cell Sci 118:3839–3847. doi:10.1242/jcs.02496

Protter DSW, Parker R. 2016. Principles and Properties of Stress Granules. Trends Cell Biol 26:668–679. doi:10.1016/j.tcb.2016.05.004

Ramírez-Peinado S, Ignashkova TI, van Raam BJ, Baumann J, Sennott EL, Gendarme M, Lindemann RK, Starnbach MN, Reiling JH. 2017. TRAPPC13 modulates autophagy and the response to Golgi stress. J Cell Sci 130:2251–2265. doi:10.1242/jcs.199521

Sacher M, Shahrzad N, Kamel H, Milev MP. 2018. TRAPPopathies: An emerging set of disorders linked to variations in the genes encoding transport protein particle (TRAPP)-associated proteins. Traffic Cph Den. doi:10.1111/tra.12615

Saleem MA, O’Hare MJ, Reiser J, Coward RJ, Inward CD, Farren T, Xing CY, Ni L, Mathieson PW, Mundel P. 2002. A conditionally immortalized human podocyte cell line demonstrating nephrin and podocin expression. J Am Soc Nephrol JASN 13:630–638.

Saurus P, Kuusela S, Dumont V, Lehtonen E, Fogarty CL, Lassenius MI, Forsblom C, Lehto M, Saleem MA, Groop P-H, Lehtonen S. 2016. Cyclin-dependent kinase 2 protects podocytes from apoptosis. Sci Rep 6:21664. doi:10.1038/srep21664

Scrivens PJ, Noueihed B, Shahrzad N, Hul S, Brunet S, Sacher M. 2011. C4orf41 and TTC-15 are mammalian TRAPP components with a role at an early stage in ER-to-Golgi trafficking. Mol Biol Cell 22:2083–2093. doi:10.1091/mbc.E10-11-0873

Sidrauski C, McGeachy AM, Ingolia NT, Walter P. 2015. The small molecule ISRIB reverses the effects of eIF2α phosphorylation on translation and stress granule assembly. eLife 4. doi:10.7554/eLife.05033

Takahashi M, Higuchi M, Matsuki H, Yoshita M, Ohsawa T, Oie M, Fujii M. 2013. Stress Granules Inhibit Apoptosis by Reducing Reactive Oxygen Species Production. Mol Cell Biol 33:815–829. doi:10.1128/MCB.00763-12

Thedieck K, Holzwarth B, Prentzell MT, Boehlke C, Kläsener K, Ruf S, Sonntag AG, Maerz L, Grellscheid S-N, Kremmer E, Nitschke R, Kuehn EW, Jonker JW, Groen AK, Reth M, Hall MN, Baumeister R. 2013. Inhibition of mTORC1 by Astrin and Stress Granules Prevents Apoptosis in Cancer Cells. Cell 154:859–874. doi:10.1016/j.cell.2013.07.031

Tiller GE, Hannig VL, Dozier D, Carrel L, Trevarthen KC, Wilcox WR, Mundlos S, Haines JL, Gedeon AK, Gecz J. 2001. A recurrent RNA-splicing mutation in the SEDL gene causes X-linked spondyloepiphyseal dysplasia tarda. Am J Hum Genet 68:1398–1407. doi:10.1086/320594

Tillmann KD, Reiterer V, Baschieri F, Hoffmann J, Millarte V, Hauser MA, Mazza A, Atias N, Legler DF, Sharan R, Weiss M, Farhan H. 2015. Regulation of Sec16 levels and dynamics links proliferation and secretion. J Cell Sci 128:670–682. doi:10.1242/jcs.157115

Tisdale EJ, Bourne JR, Khosravi-Far R, Der CJ, Balch WE. 1992. GTP-binding mutants of rab1 and rab2 are potent inhibitors of vesicular transport from the endoplasmic reticulum to the Golgi complex. J Cell Biol 119:749–761.

van Leeuwen W, van der Krift F, Rabouille C. 2018. Modulation of the secretory pathway by amino-acid starvation. J Cell Biol 217:2261–2271. doi:10.1083/jcb.201802003

Venditti R, Scanu T, Santoro M, Di Tullio G, Spaar A, Gaibisso R, Beznoussenko GV, Mironov AA, Mironov A, Zelante L, Piemontese MR, Notarangelo A, Malhotra V, Vertel BM, Wilson C, De Matteis MA. 2012. Sedlin controls the ER export of procollagen by regulating the Sar1 cycle. Science 337:1668–1672. doi:10.1126/science.1224947

Vicinanza M, Di Campli A, Polishchuk E, Santoro M, Di Tullio G, Godi A, Levtchenko E, De Leo MG, Polishchuk R, Sandoval L, Marzolo M-P, De Matteis MA. 2011. OCRL controls trafficking through early endosomes via PtdIns4,5P_2_-dependent regulation of endosomal actin. EMBO J 30:4970–4985. doi:10.1038/emboj.2011.354

Westlake CJ, Baye LM, Nachury MV, Wright KJ, Ervin KE, Phu L, Chalouni C, Beck JS, Kirkpatrick DS, Slusarski DC, Sheffield VC, Scheller RH, Jackson PK. 2011. Primary cilia membrane assembly is initiated by Rab11 and transport protein particle II (TRAPPII) complex-dependent trafficking of Rabin8 to the centrosome. Proc Natl Acad Sci U S A 108:2759–2764. doi:10.1073/pnas.1018823108

Wheeler JR, Matheny T, Jain S, Abrisch R, Parker R. n.d. Distinct stages in stress granule assembly and disassembly. eLife 5. doi:10.7554/eLife.18413

Wilson BS, Nuoffer C, Meinkoth JL, McCaffery M, Feramisco JR, Balch WE, Farquhar MG. 1994. A Rab1 mutant affecting guanine nucleotide exchange promotes disassembly of the Golgi apparatus. J Cell Biol 125:557–571.

Wippich F, Bodenmiller B, Trajkovska MG, Wanka S, Aebersold R, Pelkmans L. 2013. Dual specificity kinase DYRK3 couples stress granule condensation/dissolution to mTORC1 signaling. Cell 152:791–805. doi:10.1016/j.cell.2013.01.033

Yamasaki A, Menon S, Yu S, Barrowman J, Meerloo T, Oorschot V, Klumperman J, Satoh A, Ferro-Novick S. 2009. mTrs130 is a component of a mammalian TRAPPII complex, a Rab1 GEF that binds to COPI-coated vesicles. Mol Biol Cell 20:4205–4215. doi:10.1091/mbc.e09-05-0387

Zacharogianni M, Aguilera-Gomez A, Veenendaal T, Smout J, Rabouille C. 2014 A stress assembly that confers cell viability by preserving ERES components during amino-acid starvation. eLife 3. doi:10.7554/eLife.04132

Zou S, Liu Y, Zhang XQ, Chen Y, Ye M, Zhu X, Yang S, Lipatova Z, Liang Y, Segev N. 2012. Modular TRAPP complexes regulate intracellular protein trafficking through multiple Ypt/Rab GTPases in Saccharomyces cerevisiae. Genetics 191:451–460. doi:10.1534/genetics.112.139378

Zuscik MJ, Hilton MJ, Zhang X, Chen D, O’Keefe RJ. 2008. Regulation of chondrogenesis and chondrocyte differentiation by stress. J Clin Invest 118:429–438. doi:10.1172/JCI34174

